# Caliban: Accurate cell tracking and lineage construction in live-cell imaging experiments with deep learning

**DOI:** 10.1101/803205

**Authors:** Morgan Sarah Schwartz, Erick Moen, Geneva Miller, Tom Dougherty, Enrico Borba, Rachel Ding, William Graf, Edward Pao, David Van Valen

## Abstract

While live-cell imaging is a powerful approach to studying the dynamics of cellular systems, converting these imaging data into quantitative, single-cell records of cellular behavior has been a longstanding challenge. Deep learning methods have proven capable of performing cell segmentation—a critical task for analyzing live-cell imaging data—but their performance in cell tracking has been limited by a lack of dynamic datasets with temporally consistent single-cell labels. We bridge this gap through the integrated development of labeling and deep learning methodology. We present a new framework for scalable, human-in-the-loop labeling of live-cell imaging movies, which we use to label a large collection of movies of fluorescently labeled cell nuclei. We use these data to create a new deep-learning-based cell-tracking method that achieves state-of-the-art performance in cell tracking. We have made all of the data, code, and software publicly available with permissive open-source licensing through the DeepCell project’s web portal https://deepcell.org.

---

Live-cell imaging, in which cells are imaged over time with light microscopy, provides a window into the dynamic behavior of living cells. Data generated by this class of experiments have shed light on numerous cellular processes, including cellular heterogeneity^1–4^, cell division^5,6^, morphological transitions^7–11^, and signal transduction^12–15^. New technologies that pair perturbations with imaging have led to renewed interest in using imaging to phenotype cellular dynamics^16–20^. While powerful, live-cell imaging data present a significant challenge to rigorous quantitative analysis. Central to the analysis of these data is single-cell analysis, where each cell is detected and tracked over time. Accurate solutions to these two tasks—cell detection and tracking—are essential components of every live-cell imaging analysis pipeline.

Modern deep learning methods offer a compelling path to general solutions to the computer vision problems raised by cellular imaging data. While powerful, the performance of these methods is limited by the availability of labeled data that pairs example images with information-rich labels. Researchers have made substantial progress in cell segmentation, primarily because of the increased availability of labeled data and the development of human-in-the-loop (HITL) labeling methodology for static images^21–23^. Progress in deep learning solutions to cell tracking has been more limited due to a lack of similar data resources and methodology for dynamic data. Existing datasets (Table 1) are limited in their scope and scale^24–31^, whereas simulated datasets have not yet proven capable of creating high-performing models^24,32,33^. Further, existing datasets are limited in the resolution of their labels (e.g., point labels vs. pixel-level segmentation labels), trajectory length (the number of frames over which a cell is tracked), and the number of mitotic events (Table 1). These limitations are understandable, given the time-consuming nature of labeling dynamic movies. Not only must each cell be segmented in a temporally consistent way, but lineage information must also be captured by tracking cells over time and labeling cell division events. Existing labeling methodology that has proven scalable for static images has yet to be extended at scale to these dynamic datasets^34^. In this work, we applied a full-stack approach to the problem of cell tracking, with a specific focus on tracking fluorescently labeled cell nuclei in mammalian cells. Specifically, we combined a HITL approach to image labeling^22^ adapted to dynamic imaging data, a novel deep learning algorithm for cell tracking, and new benchmarks for cell tracking to create a new labeled reference dataset for cell tracking. We used this dataset—DynamicNuclearNet—to develop state-of-the-art deep learning models for cell tracking. We further integrated these models into a pipeline called Caliban, which enables rapid and accurate segmentation, tracking, and lineage construction of nuclear live-cell imaging data with no manual parameter tuning. The source code described in this work is available at https://github.com/vanvalenlab/deepcell-tf and https://github.com/vanvalenlab/deepcell-tracking; datasets and pre-trained models are available through our lab’s web portal https://deepcell.org.

**Table 1:** Publicly available labeled datasets for two-dimensional temporal (2D+T) cell tracking. CTMC: Cell Tracking with Mitosis Detection Dataset Challenge, CTC: Cell Tracking Challenge, Fluor: fluorescent, DIC: differential interference contrast, WC: whole cell, Part: partial.

| Name | Modality | Annotation Type | Cell Types | Objects | Tracks | Divisions | Source |
| --- | --- | --- | --- | --- | --- | --- | --- |
| DynamicNuclearNet | Fluor. Nuclei | Nuclear Mask | 5 | 647,322 | 16,501 | 2,621 | This work |
| DeepSea | Phase | Nuclear Mask | 3 | 100,000 | 2,576 | 137 | Zargari *et al.*<sup>31</sup> |
| BTrack/CellX | Fluor. Nuclei | Nuclear Mask | 1 | - | 1,032 | - | Cuny *et al.*<sup>36</sup> |
| CTMC | DIC | Bounding Box | 14 | 2,045,834 | 2,900 | 457 | Anjum & Gurari<sup>27</sup> |
| C2C12 | Phase | Centroid | 1 | 135,859 | 23,400 | 7,159 | Ker *et al.*<sup>25</sup> |
| CTC 2D+T | DIC | Part. WC Mask | 1 | - | 70 | 18 | CTC<sup>37</sup> |
| CTC 2D+T | Phase | Part. WC Mask | 2 | - | 2,415 | 1,019 | CTC<sup>37</sup> |
| CTC 2D+T | Fluor. Nuclei | Part. Nuclear Mask | 3 | - | 1,227 | 395 | CTC<sup>37</sup> |
| CTC 2D+T | Fluor. WC | Part. WC Mask | 2 | - | 128 | 10 | CTC<sup>37</sup> |
| CTC 2D+T | Brightfield | Part. WC Mask | 2 | - | 459 | 242 | CTC<sup>37</sup> |

## 1 Results

### 1.1 HITL labeling of large-scale, dynamic imaging data

We employed two key strategies to label dynamic imaging data efficiently. First, we made use of crowdsourcing to parallelize our work. Second, we utilized a HITL approach to accelerate labeling efforts. Our approach had two phases: cell segmentations were generated in the first phase, whereas cells were tracked and cell divisions were labeled in the second phase (Fig. 1a). The segmentation phase of our approach followed prior work^22^, beginning with a small seed dataset for cell segmentation generated by expert labelers. We then trained a preliminary model to generate candidate segmentations, which were refined through a round of crowdsourced correction and expert quality control (QC). We used DeepCell Label, our browser-based labeling engine specifically designed for cellular images for this work. Critically, labelers were shown a sequence of five frames rather than individual frames to leverage temporal information to increase the label accuracy and to ensure temporal consistency before labeling cell lineages. After a sufficient amount of data was labeled, the model was retrained on a new dataset that combined the original seed dataset and the corrected predictions. The updated model was then used to generate subsequent segmentation labels. This cycle was repeated until model predictions matched expert predictions, as judged by qualitative comparison and quantitative metrics (Section 3.5.5). In the tracking phase, labelers were given a complete movie (42-92 frames) and tasked with tracking cells and identifying cell divisions. This phase was achieved through iterative cycles of model prediction, crowdsourced correction, expert QC, and model retraining. For this task, we extended DeepCell Label to include tools for labeling cell lineages and divisions. To coordinate this multi-stage dataset development process, we implemented a data and model versioning system using Data Version Control (DVC)^35^, which acts alongside Git to track each data file and its associated metadata (Fig. 1a). By automating these file associations, we removed the need for an expert user to manage the labeling pipeline and keep track of various computational notebooks or scripts.

**Figure 1:**
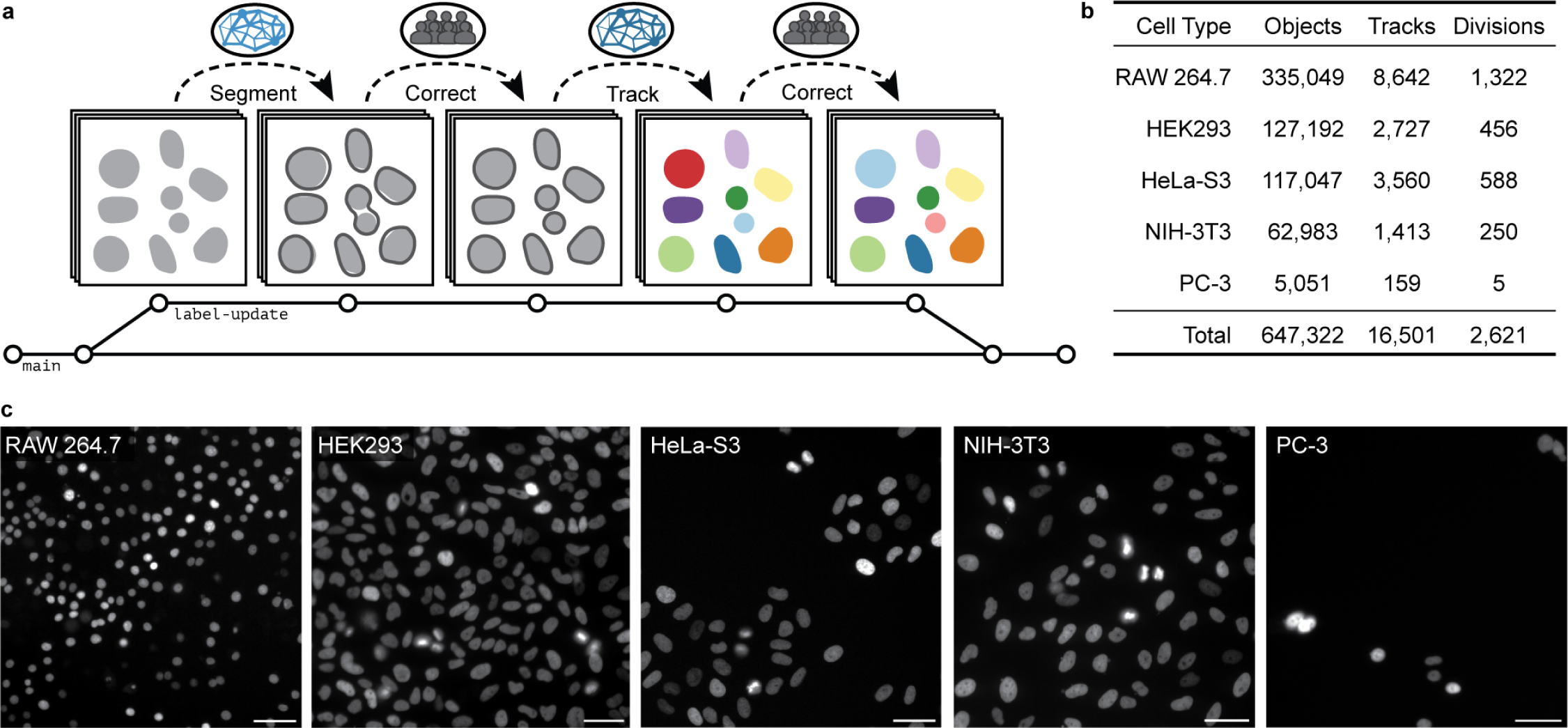
Developing DynamicNuclearNet with HITL annotation of dynamic data. (a) Our HITL process for generating labels alternated between preliminary models generating predictions and human annotators correcting errors generated by the model. This process was conducted twice: first for segmentation and again for tracking. Each update to the labeled data was versioned and saved with DVC^35^. (b) With a two-stage HITL process, we assembled DynamicNuclearNet, a dataset of segmented and tracked dynamic cell nuclei encompassing five cell lines. (c) Example images of each of the five cell lines. (Scale bar = 50 *µ*m)

Using this methodology, we built DynamicNuclearNet, a segmented and tracked dataset of fluorescently labeled cell nuclei spanning five different cell lines. This dataset contains 647,322 unique nuclear segmentations assembled into over 16,501 trajectories with over 2,621 division events (Fig. 1b,c). Each trajectory begins at the cell’s appearance in the field of view (FOV) or birth as a daughter cell and ends when the cell disappears by permanently leaving the FOV, dying or dividing. While generating pixel-level masks for each cell is expensive compared to other types of labels (e.g., centroids or bounding boxes), these masks facilitate numerous downstream analysis steps, such as quantifying signaling reporters or nuclear morphology. The 2,621 division events in our dataset surpass all previous annotation efforts that utilize nuclear segmentation masks (Table 1), which allows us to incorporate cell division detection into our deep-learning-based cell-tracking method.

### 1.2 Accurate nuclear segmentation and tracking with Caliban

In tandem with the labeling methodology advances described above, we developed Caliban, an integrated solution to nuclear segmentation and tracking. Caliban employs a tracking-by-detection approach in which cells are first identified in each frame by a deep learning model; these detections are then used to reconstruct a lineage tree that connects cells across frames and through cell division events. For the reconstruction of lineage trees, we utilize a deep learning model inspired by Sadeghian *et al.*^41^, which encodes temporal dependencies for multiple features of each object and predicts the probability of a parent– child relationship that exists due to a cell division event between any pair of cells across frames. In this approach, accurate cell detection and segmentation are essential to producing faithful lineage reconstructions. To this end, we have combined our prior work on cell segmentation^22,42^ with DynamicNuclearNet and a comprehensive benchmarking framework to train an accurate deep learning model for nuclear segmentation as part of Caliban.

The processing steps for Caliban are shown in Fig. 2a. Raw images are passed through the nuclear segmentation model to produce cell masks. These masks are used to extract features for each cell, while the centroids are used to construct an adjacency matrix to identify cells in close proximity (*<* 64 pixels, 41.6 *µ*m). These features and the adjacency matrix are fed into a neighborhood encoder model, which uses a graph attention network^38,39^ to generate feature vectors that summarize information about a cell’s—and its neighbors’—appearance, location, and morphology (Fig. 2b). These feature vectors are then fed into a tracking model that causally integrates temporal information and performs a pairwise comparison of each cell’s feature vector across frames to produce an effective probability score indicating whether two cells are the same cell, are different cells, or have a parent–child relationship (Fig. 2c). Separating our tracking model into two pieces also facilitates rapid and scalable inference. During inference, the computationally expensive neighborhood encoder model can be run on all frames in parallel, leveraging GPU acceleration, followed by the lightweight tracking inference model, which is run on a frame-by-frame basis. The tracking inference model assigns lineages to cells by comparing the feature vectors of the last frame of existing lineages with the feature vectors of candidate cells in the current frame; model predictions are used with the Hungarian algorithm^43,44^ to complete the assignment. To accommodate the entry and exit of cells in the linear assignment framework, we create a “shadow object” for each cell in the frame, which allows assignments for the “birth” or “death” of cells in each frame^44^. The methods (Section 3.5) provide full details of the model architecture, model training, postprocessing, and hyperparameter optimization.

**Figure 2:**
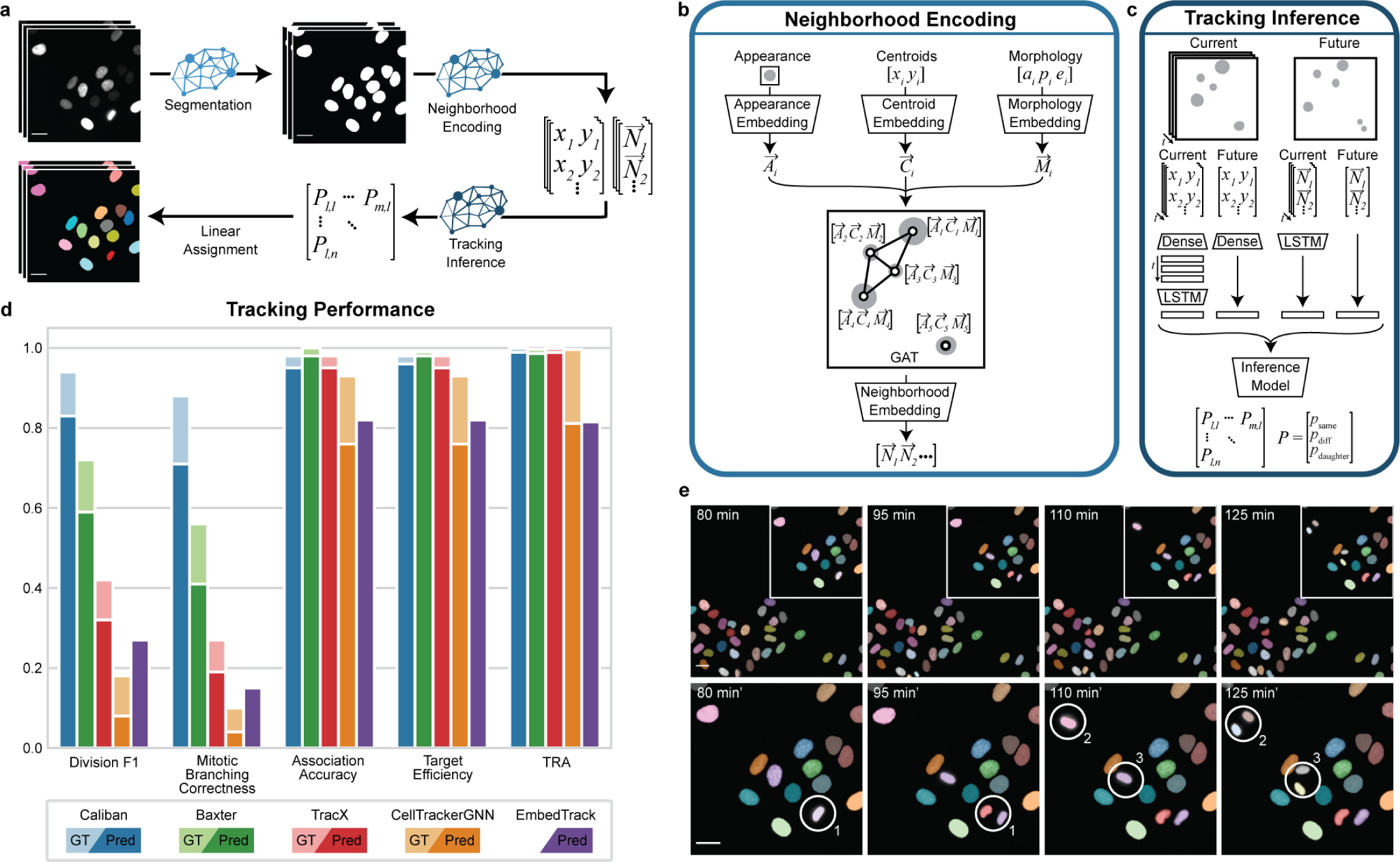
A deep learning approach to cell segmentation and tracking using Caliban. (a) Caliban takes a movie of fluorescently labeled nuclei as input and then generates a nuclear segmentation mask for each frame. Features for each cell in a frame are extracted and passed through a neighborhood encoder model to generate a vector embedding for each cell. These embeddings and cell positions are passed into the tracking inference model, which predicts the probability that each pair of cells between frames is the same, is different, or has a parent–child relationship. These probabilities are used as weights for linear assignment to construct cell lineages on a frame-by-frame basis. (b) The neighborhood encoding model takes as input an image of each cell, its centroid position, and three metrics of morphology (area, perimeter, and eccentricity). A vector embedding of each input is used as node weights in a graph attention network^38,39^, where edges are assigned to cells within 64 pixels (41.6 *µ*m) of each other. The final neighborhood embedding for each cell captures the appearance of that cell and its spatial relationship with its neighbors in that frame. (c) The tracking inference model performs pairwise predictions on cells in frame *t_n_* to cells in frame *t_n_*_+1_. The model is given neighborhood embeddings and centroid positions of cells in the previous seven frames [*t_n−_*_7_*, t_n_*] to compare with cells in frame *t_n_*_+1_. The temporal context of the previous seven frames is modeled using long short-term memory (LSTM) layers^40^. Ultimately, the model outputs a set of effective probabilities (*p*_same_, *p*_diff_, and *p*_parent-child_) for each pair of cells between frame *t_n_* and frame *t_n_*_+1_. (d) The performance of Caliban and that of four other tracking methods were evaluated on the test split of DynamicNuclearNet. Tracking performance on ground truth segmentations is excluded for EmbedTrack because it is an end-to-end method that generates segmentations as part of tracking. TRA: tracking accuracy in the Cell Tracking Challenge. (e) A sample montage from DynamicNuclearNet with predictions from Caliban. Circles highlight the correct identification of three division events. (Scale bars = 26 *µ*m)

In addition to Caliban, we also developed a comprehensive framework for evaluating tracking performance as part of this work. Most existing metrics for cell tracking focus on the quality of linkages between cells in lineage trees^45–48^. While useful, these approaches mask the method’s performance on cell division. Many downstream analyses rely on accurate division detection; however, the relative rarity of division events makes them difficult to assess with summary metrics. To resolve this issue, we implemented several existing evaluation metrics that quantify a method’s performance on cell division^30,49,50^; the details of each metric are given in the methods (Section 3.5.5). Our open-source Python package allows these benchmarks to be computed for any dataset saved in the standard CTC format.

We used these metrics in combination with prior metrics^30,45–47^ to compare Caliban against four alternative algorithms for cell tracking: Baxter, CellTrackerGNN, EmbedTrack, and Trac^x^. We focused on methods that performed well in the Cell Tracking Challenge and could be run without manual parameter tuning – including using parameters published as part of the challenge submission. Baxter implements the Viterbi algorithm^51^. CellTrackerGNN constructs a global track solution by pairing a graph neural network with an edge classifier to extract cell lineages^52^. EmbedTrack utilizes a single convolutional neural network for joint cell segmentation and tracking^53^. Trac^x^ pairs a feature-based linear assignment problem with a cell fingerprint classifier to curate tracking results^36^. For each of these methods, we utilized pre-existing models or parameters that were trained or optimized on the Fluo-N2DL-HeLa dataset if available. For Caliban, we used a single version of the model that was trained on all five cell types represented in DynamicNuclearNet, including the two Fluo-N2DL-HeLa training movies. We tested each algorithm on ground truth and predicted segmentations. Predicted segmentations for each method were generated with that method’s segmentation model or Caliban’s segmentation model if the former was unavailable. On measures of division performance evaluated on the DynamicNuclearNet testing split, Caliban outperformed all previously published methods. This performance boost is primarily attributable to Caliban’s cell-tracking capability rather than cell segmentation (Fig. 2e), as the performance boost is present when tracking is performed on ground truth segmentations. On metrics focused on segmentation and linkages evaluated on the DynamicNuclearNet testing split, Caliban performed comparably to existing methods. For all metrics evaluated on the Cell Tracking Challenge Fluo-N2DL-HeLa test split, Caliban outperformed previously published methods. Complete benchmarking results are shown in Supplementary Tables 1 and 2. We note that these benchmarks are unable to separate the relative contributions of training data size and model architecture to performance.

To increase the accessibility of Caliban, we have made Caliban available through our lab’s GitHub (https://github.com/vanvalenlab/deepcell-tf) and the DeepCell web portal (https://deepcell.org). Converting algorithms into robust, user-friendly software can be challenging, particularly for cell-tracking algorithms, given the large size and varied nature of the data on which these algorithms must operate^24,32,34,54^. Here, we leverage our prior work in developing the DeepCell Kiosk^55^, a cloud-native, scalable software platform for cellular image analysis pipelines that use deep learning methods. Our cloud deployment of Caliban was facilitated by our emphasis on inference speed, scalability, and accuracy during method development. We believe that the availability of Caliban in both local and cloud versions will make this tool accessible to the broader life science community. Further, the open-source datasets, models, and benchmarking tools will facilitate future method development.

## 2 Discussion

Live-cell imaging is a transformative technology critical to elucidating cellular proliferation, migration, and other dynamic phenomena. The utility of this technology has long been limited by our ability to extract quantitative, single-cell information from these movies. In this work, we have made a significant step toward solving the computer vision challenges of dynamic cellular imaging data. By extending scalable, HITL labeling frameworks^21,22,56–59^ to dynamic data, we have generated a labeled dataset of over 16,000 cellular trajectories and 2,600 cell divisions. We demonstrated that these labeled data can power accurate nuclear segmentation and tracking models. We further showed that these models can be combined into an integrated pipeline and deployed on a cloud-native platform for scalable, user-friendly inference.

While we achieved impressive performance on nuclear segmentation and tracking, several areas of improvement exist for future work. First, accurate cell segmentation remains a performance bottleneck, as highlighted by the difference in the performance of tracking methods on ground truth and deep-learning-generated segmentations (Fig. 2d). Additional performance gains on segmentation will likely arise via methods that leverage temporal information to improve performance. Newer segmentation methods that enable the segmentation of overlapping objects^23,60^—a limitation of all cell segmentation methods that currently see wide use^61^—may also help close this gap. Second, while the dataset we have collected is impressive in scale, its diversity remains limited compared with the full space of live-cell phenotypes. Our focus on cell nuclei allowed us to develop a new labeling methodology for dynamic datasets; creating a dataset similar in label scale and quality for whole cells will likely be the focus of future work. Such work may leverage recent methods like the Segment Anything Model from Meta to further accelerate labeling throughput^62,63^. To aid this effort, we have already extended the features available in DeepCell Label to support the annotation and tracking of overlapping cells. Moreover, dynamic cell phenotypes change substantially in the setting of perturbations. Such shifts in data distribution are expected to degrade cell segmentation and tracking performance; the best approach for mitigating this issue is to expand the space of labeled data to capture these phenotypes. Doing so will require expanding our labeling framework to capture more dynamic phenotypes (e.g., cell death). We note that imaging data from pooled optical screens^16,18,20,64,65^ may also be a valuable path for generating images of perturbed cell phenotypes at scale. Third, our work focuses on 2D live-cell imaging movies. While much information can be extracted from this form of data, their 3D counterparts are of high value to many life science communities. Extending the methodology presented here to 3D movies should be the focus of future work.

Our work contains several lessons for the community of researchers developing deep-learning methods for cellular image analysis. First, our work highlights the importance of data labeling methodology and data scale. By developing a scalable approach for labeling dynamic live-cell imaging data, we have compiled a pixel-level labeled dataset that is substantially larger than previous datasets. The increased scale of the data allowed us to compile enough examples of cell divisions—a critical but rare dynamic event—to enable accurate detection by deep learning models. While models trained on sparsely labeled data can be effective^34,50^, increasing the amount of labeled data is essential to creating models that generalize across datasets. Second, our study demonstrates the importance of informative benchmarks. As highlighted by our work and that of others^30,49,50^, accurate cell division detection is one of the most challenging aspects of cell tracking but is critical to constructing cell lineages. This task is challenging for supervised methods, largely because of the class imbalance—cell divisions represent a relatively minor fraction of linkages in cell lineage trees. Aggregate metrics for cell tracking mask cell division events, making it difficult to judge performance gains during and after algorithmic method development^45^. This limitation is evident when comparing the CTC’s TRA metric to metrics that are specific to cell division. Methods that perform well on the TRA score can have poor performance on the cell division task. Combining aggregate metrics with specific, informative metrics creates a more complete picture of performance and is critical to crafting methods that can be used in production. Finally, this work underscores the value of model scalability. While accuracy is often the metric used to judge cellular image analysis methods, inference speed—and hence scalability—is equally important. Faster workflows can process substantially more data and provide a better user experience. A major focus of this work was scalability, which was achieved by crafting an architecture in which computationally expensive operations (e.g., the neighborhood encoder) were performed in parallel on specialized hardware (e.g., GPUs). Our model’s scalability enables a responsive cloud deployment—a typical dataset (10,000 cell detections over 30 frames) can be processed in *∼* 40 s on an A6000 GPU, with much of the processing time taken by segmentation (Supplementary Fig. 1).

In conclusion, our work provides the live-cell imaging community with an accessible, accurate method for reconstructing single-cell lineages from dynamic imaging experiments. We believe our work will facilitate the analysis of cell phenotypes and behaviors in a wide variety of high-throughput imaging experiments.

## 3 Materials and Methods

### 3.1 Cell culture

We used five mammalian cell lines (NIH-3T3, HeLa-S3, HEK293, RAW 264.7, and PC-3) to collect training data. All lines were acquired from the American Type Culture Collection. We cultured the cells in Dulbecco’s modified Eagle’s medium (DMEM; Invitrogen; RAW 264.7, HEK293, and NIH-3T3) or F-12K medium (Caisson; Hela-S3 and PC-3) supplemented with 2 mM L-glutamine (Gibco), 100 U/mL penicillin, 100 *µ*g/ml streptomycin (Gibco), and either 10% calf serum (Colorado Serum Company) for NIH-3T3 cells or 10% fetal bovine serum (FBS; Gibco) for all other cells.

### 3.2 Live imaging

Before imaging, cells were seeded in fibronectin-coated (10 *µ*g/mL; Gibco) glass 96-well plates (Nunc or Cellvis) and allowed to attach overnight. We performed nuclear labeling via prior transduction with H2B-iRFP670 (Hela, RAW 264.7), H2B-mClover (HEK293, NIH/3T3), and H2B-mCherry (PC-3). The media was removed and replaced with imaging media (FluoroBrite DMEM (Invitrogen) supplemented with 10 mM HEPES (Sigma-Aldrich), 10% FBS (Gibco), 2mM L-glutamine (Gibco)) at least 1 h before imaging. We imaged cells with a Nikon Ti-E or Nikon Ti2 fluorescence microscope with environmental control (37*^◦^*C, 5% CO_2_) and controlled by Micro-Manager or Nikon Elements. We acquired images at 5- to 6-min intervals with a 20x objective (40x for RAW 264.7 cells) and either an Andor Neo 5.5 CMOS camera with 3*×*3 binning or a Photometrics Prime 95B CMOS camera with 2*×*2 binning. All data were scaled so that pixels had the same physical dimensions (0.65 *µ*m per pixel) before training.

### 3.3 Dataset development

#### 3.3.1 DeepCell Label

We previously described DeepCell Label^22^, our browser-based software for data annotation. We extended DeepCell Label to support labeling cell lineages and divisions in dynamic datasets. Additionally, we implemented a state machine that allows annotators to apply undo/redo functions during their work. These new features are described below.

DeepCell Label manages the state of its labeled data with a Python-based backend and a React-based frontend. The backend serves and submits project data and provides image processing for editing label arrays, while the frontend controls the user interface and edits non-image-based labels. The front end retrieves images and labels for the project from the backend and loads the data into its state management.

We use the Javascript library XState to manage the state on the front end. Through XState, we define actors that manage the state of both user interface (UI) elements and data labels. An actor consists of a context, containing arbitrary data, along with a set of states and transitions between states. Actors receive events from other actors or the user, and each state defines how to transition between states and update its context upon receiving events. Some actors maintain UI state, while others define and control the operations that can edit labeled data. The application creates a root actor that instantiates a tree of child actors corresponding to each UI element or type of labeled data. The root actor sets up communication between child actors, enabling features like undo/redo that must orchestrate state across the application.

Actors expose their state to user interface elements by React’s hooks, allowing multiple components to access and update the same shared state. For instance, to adjust the contrast of an image, a Slider can expose the contrast settings for a user to update, while a Canvas component can access the updated contrast settings to render the image. For example, we define an actor for the image canvas that handles mouse movement events with context-dependent behavior. When panning around the canvas, a mouse-move event changes the position of the canvas, while when not panning, we update the coordinates for the cursor. Clicking and releasing the mouse sends mouse-up and mouse-down events, transitioning to and from the panning state.

We implement our undo feature via an undo actor that records and restores the states of both UI and label actors over time. The undo actor maintains two stacks of project states: an undo stack with past snapshots to restore and a redo stack with undone snapshots. Editing the labels after undoing clears the redo stack, so only one branch of the project state is maintained. To integrate with the undo feature, actors register themselves with the undo actor and agree to submit snapshots of their state that can be restored upon undoing or redoing an action. Two types of actors can register with the undo actor: UI actors that manage the state of UI elements, and label actors that manage and edit labels. When a label actor edits its labels, it submits a SNAPSHOT event to the undo actor containing a copy of the labels before and after editing. The undo machine then broadcasts a SAVE event to all UI actors, which respond with a RESTORE event containing their current state. When the user undoes an action, the SNAPSHOT and RESTORE events are resent to the actors that originally submitted the events. With this approach, each UI and label actor is responsible for defining how its state should be recorded and restored with the events it submits, and the undo actor is responsible only for orchestrating and broadcasting the events to all registered actors. As new types of labeled data and UI elements are developed, the actors that drive new features can flexibly integrate with the undo infrastructure by defining these events for themselves and registering with the undo actor.

#### 3.3.2 Data annotation

In this study, we utilized DeepCell Label in two stages to generate a nuclear tracking dataset. First, annotators were asked to correct nuclear segmentation labels for all frames in the dataset. Movies were broken into five frame sets for segmentation which allows annotators to leverage the temporal context present in the movie to improve the annotation of dividing cells. Second, after segmentation annotations were complete, annotators were asked to label the nuclear segmentation masks such that a single cell maintains the same label across frames. Additionally, all division events were annotated with the connection of the parent cell to each daughter cell. An expert annotator reviewed all annotated movies before incorporation into the training dataset. The supplementary information provides a user manual for DeepCell Label (Supplementary File 1), along with sample instructions for segmentation (Supplementary File 2) and tracking (Supplementary File 3) corrections. Annotations were conducted by a team of four annotators and two expert reviewers. Each movie was annotated by a single annotator and approved by a single expert eliminating the need for resolving differences between two independent annotators.

#### 3.3.3 Data versioning with DVC

Each labeled movie was versioned and tracked with DVC^35^. We recorded additional metadata in each .dvc file, including the data dimensions, annotation progress, and data source. These metadata enabled automatic data processing for generating segmentation and tracking predictions as well as launching annotation tasks.

#### 3.3.4 Dataset sources

DynamicNuclearNet contains data from six sources. Five datasets were collected internally as described in sections 3.1 and 3.2. Additionally, the CTC dataset Fluo-N2DL-HeLa was incorporated after generating complete segmentation masks for all frames using the protocol described above.

### 3.4 Nuclear segmentation

#### 3.4.1 Deep learning model architecture

The deep learning model for nuclear segmentation was based on the design of feature pyramid networks^66,67^. The network was constructed from an EfficientNetV2L backbone^68^ connected to a feature pyramid. Input images were concatenated with a coordinate map before entering the backbone. We used backbone layers C1–C5 and pyramid layers P1–P7. The final pyramid layers were connected to three semantic segmentation heads that predict transforms of the labeled image.

#### 3.4.2 Label image transforms

For each image, we used a deep learning model to predict three different transforms, as inspired by previous work^22,69,70^. The first transform predicted whether a pixel belongs to the foreground or background, known as the “foreground–background transform.” The second transform predicted the distance of each pixel in a cell to the center of the cell and is called the “inner distance.” If the distance between a pixel and the center of the cell is *r*, then we compute the transform as 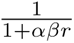, where 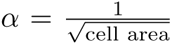 and *β* is a hyperparameter set to 1^22^. The final transform was the “outer distance,” which is the Euclidean distance transform of the labeled image. The loss function was computed as the sum of the mean squared error on the inner and outer distance transforms and the weighted categorical cross-entropy^71^ on the foreground–background transform. The cross-entropy term was scaled by 0.01 before the sum.

#### 3.4.3 Preprocessing

Each image was required to have a minimum of one labeled object. Additionally, each image was normalized using contrast-limited adaptive histogram equalization with a kernel size equal to 1/8 of the image size to ensure that all images have the same dynamic range^72^.

#### 3.4.4 Postprocessing

We fed two of the three model outputs, the inner and outer distance, into a marker-based watershed method^73^ to convert the continuous model outputs into a discrete labeled image in which each cell is assigned a unique integer. We applied a peak-finding algorithm^74^ with a radius of 10 pixels and a threshold of 0.1 to the inner distance prediction to determine the centroid location of each cell. Next, we generated the cell mask image by applying the watershed algorithm to the inverse outer distance prediction with the centroids as markers and a threshold of 0.01.

#### 3.4.5 Model training and optimization

Training data were augmented with random rotations, crops, flips, and scaling to improve the diversity of the data. We used 70% of the data for training, 20% for validation, and 10% for testing. The model was trained using the Adam optimizer^75^ with a learning rate of 10*^−^*^4^, a clipnorm of 10*^−^*^3^, and a batch size of sixteen images; training was performed for sixteen epochs. After each epoch, the learning rate was adjusted using the function lr = lr *×* 0.99^epoch^. Additionally, if the loss of the validation data did not improve by more than 10*^−^*^3^ after five epochs, the learning rate was reduced by a factor of 0.01.

To optimize the model’s performance on nuclear segmentation, we tested ten backbones: ResNet50^76^, ResNet101^76^, EfficientNetB2^77^, EfficientNetB3^77^, EfficientNetB4^77^, EfficientNetV2M^68^, EfficientNetV2L^68^, EfficientNetV2B1^68^, Efficient-NetV2B2^68^, and EfficientNetV2B3^68^. Additionally, we explored the optimal set of pyramid layers: P1–P7 and P2–P7.

#### 3.4.6 Evaluation

To fully evaluate the performance of our segmentation model, we developed a set of object-based error classes that assess the model on a per-object basis as opposed to a per-pixel basis. This framework provided a perspective on model performance that reflects downstream applications. First, we built a cost matrix between cells in the ground truth and cells in the prediction, where the cost is one minus the intersection over union (IoU) for each pair of cells. We performed a linear sum assignment on this cost matrix, with a cost of 0.4 for unassigned cells, to determine which cells were correctly matched between the ground truth and prediction. For all remaining cells, we constructed a graph in which an edge was established between a ground truth and a predicted cell if the IoU was greater than zero. For each subgraph, we classified the error type based on the connectivity of the graph. Nodes without edges corresponded to a false positive or negative if the graph contained only a predicted or ground truth cell, respectively (Supplementary Fig. 2a–c). A single predicted node connected to multiple ground truth nodes indicated a merge error (Supplementary Fig. 2d). Conversely, a single ground truth node connected to multiple predicted nodes was a split error (Supplementary Fig. 2e). Finally, any subgraphs that contain multiple ground truth and predicted nodes were categorized as “catastrophe” (Supplementary Fig. 2f). The resulting error classes can be used to calculate a set of summary statistics, including recall, precision, and F1 score by using the true positive, false positive, and false negative classes. The remaining error classes can be used to calculate (1) the number of missed detections resulting from a merge, (2) the number of gained detections resulting from a split, (3) the number of true detections involved in a catastrophe, and (4) the number of predicted detections involved in a catastrophe.

### 3.5 Cell tracking

#### 3.5.1 Linear assignment for tracking

Our tracking algorithm drew inspiration from Jaqaman *et al.*^44^, where tracking was treated as a linear assignment problem. To solve the tracking problem, we first constructed a cost function for possible pairings across frames. The tracking problem was then reduced to the selection of one assignment out of the set of all possible assignments that minimized the cost function. This task can be accomplished with the Hungarian algorithm^78^. One complicating factor of biological object tracking is that objects can appear and disappear, which leads to an unbalanced assignment problem. Cells can disappear by either moving out of the FOV or dying. Similarly, cells can appear by moving into the FOV or dividing into two daughter cells from one parent cell. In the context of the linear assignment problem, one can preserve the runtime and performance by introducing a “shadow object” for each object in the two frames that represents an opportunity for objects to “disappear” (if an object in frame *t_n_* is matched with its shadow object in frame *t_n_*_+1_) or “appear” (if an object in frame *t_n_*_+1_ is matched with its shadow object in frame *t_n_*)^44^. Assuming that mitotic events can be accommodated by a “shadow object” as well, division detection and assignment fit neatly into this framework. This framework can also accommodate cells that disappear from the field of view and reappear, by allowing unmatched cells from prior frames that were not assigned to cell division events to participate in the assignment. With the annotated trajectories and divisions from our dataset, it then becomes a matter of developing a deep learning architecture to extract an object’s features and learn an optimal cost function.

To construct our learned cost function, we cast it as a classification task. Let us suppose that we have two cells: our target cell *i* in frame *t_n_* and cell *j* in frame *t_n_*_+1_. Our goal was to train a classifier that takes in information about each cell and produces an effective probability indicating whether these two instances are the same, are different or have a parent–child relationship. If we have already tracked several frames, we incorporate temporal information by using multiple frames of information for cell *i* as an input to the classifier. This approach allowed us access to temporal information beyond just the two frames we are comparing. Our classifier was a hybrid deep learning model that blends recurrent, convolutional, and graph submodels; its architecture is summarized in Fig. 2b,c. The three scores that the model outputs, (*p*_same_, *p*_diff_, and *p*_parent-child_), which are all positive and sum to unity, can be thought of as probabilities. These scores were used to construct the cost matrix. If a cell in frame *t_n_*_+1_ is assigned to a shadow cell, i.e., if it “appears,” then we assessed whether there is a parent–child relationship. This was done by finding the highest *p*_parent-child_ among all eligible cells (i.e., the cells in frame *t_n_* that were assigned to “disappear”)—if this probability was above a threshold, then we made the lineage assignment.

#### 3.5.2 Neighborhood encoder architecture

To capture the contextual information of each cell and its neighbors, we constructed a graph attention network^38,39^. There were three input heads to the model. The first head received images of each cell and converted these images to a vector embedding with a convolutional neural network. Each image consisted of a 16*×*16 crop of the raw data centered on the centroid position of the cell. Additionally, the pixels within the nuclear segmentation mask were normalized by subtracting the mean value and dividing by the standard deviation. The second head received the centroid location of each cell. The third head received three morphology metrics for each cell: area, perimeter, and eccentricity. The latter two heads made use of fully connected neural networks to convert the inputs into vectors. We built an adjacency matrix for the graph attention network based on the Euclidean distance between pairs of cells; cells were linked if they were closer than 64 pixels (41.6 *µ*m). The normalized adjacency matrix and concatenated embeddings were fed into a graph attention layer^38^ to update the embeddings for each cell. The appearance and morphology embeddings were concatenated to the output of the graph attention layer to generate the final neighborhood embedding.

#### 3.5.3 Tracking model architecture

Given cell 1 in frame *t_n_* and cell 2 in frame *t_n_*_+1_, the neighborhood encoder was used to generate embeddings for cell 1 in frame *t_n_* and the previous seven frames [*t_n−_*_7_*, t_n_*]. Long short-term memory^40^ layers were applied to the resulting embedding for cell 1 to merge the temporal information and to create a final summary vector for cell 1. The neighborhood encoder then generated an embedding for cell 2. Next, the vectors for cell 1 and cell 2 were concatenated and fed into fully connected layers. The final layer applied the softmax transform to produce the final classification scores: *p*_same_, *p*_diff_, and *p*_parent-child_.

#### 3.5.4 Training and optimization

Both the neighborhood encoder and the inference model were jointly trained end-to-end such that the neighborhood embedding was tuned for the inference task. The model was trained on data that compare a set of frames [*t_n−_*_7_*, t_n_*] with frame *t_n_*_+1_. Each comparison of *t_n_* with *t_n_*_+1_ contributed to the loss. For inference, the model was given single pairs of frames, e.g., *t_n_* vs. *t_n_*_+1_. Training data were augmented with random rotations and translations. We used 70% of the data for training, 20% for validation, and 10% for testing. Data splitting was performed with regard to the cell type such that each cell type is equally represented across the three splits. The model was trained using the rectified Adam optimizer^79^ with a learning rate of 10*^−^*^3^, a clipnorm of 10*^−^*^3^, and a batch size of eight images. After each epoch, the learning rate was adjusted using the function lr = lr *×* 0.99^epoch^. Additionally, if the loss of the validation data did not improve by more than 10*^−^*^4^ after five epochs, the learning rate was reduced by a factor of 0.1. The model was trained over 50 epochs.

To optimize the performance of the tracking model, we tested the following parameters: graph layers (graph convolution layer, graph convolution layer with trainable skip connections, and graph attention convolution layer), distance threshold (64, 128, 256 pixels; 41.6, 83.2, 166.4 *µ*m), crop mode (fixed and resized), birth probability, division probability, and death probability.

#### 3.5.5 Evaluating tracking performance

To evaluate the tracking performance, we utilized two sets of metrics. The first set assessed the linkages between cells, whereas the second set focused on the linkages of dividing cells. For the first set of metrics, we calculated the target efficiency (TE) and association accuracy (AA)^46,47^. Briefly, TE assesses the fraction of cells assigned to the correct lineage, and AA measures the number of correct linkages generated between cells.

Traditional metrics for evaluating tracking, including TE and AA, do not accurately reflect the ability of the method to identify divisions because divisions are relatively rare events. To overcome this weakness, we developed an evaluation pipeline that classifies each division event as a correct, missed, or incorrect division. Our pipeline can handle tracking assignments on ground truth and predicted segmentations. First, we calculated the IoU between cells in the ground truth and the predictions to establish a mapping that can be used to compare tracking predictions. On predicted segmentations, an IoU threshold of 0.6 was used as a threshold for overlap. For each division in the ground truth, we checked the corresponding node in the prediction to determine whether it was labeled as a division. If the daughter nodes in the prediction match those in the ground truth, the division was counted as a correct division (Supplementary Fig. 3a). We have found that depending on the predicted segmentations, a division can sometimes be assigned to the frame before or after the frame that is annotated as a division in the ground truth data. We treated these shifted divisions as correct or a true positive. If the predicted node was not labeled as a division, it was considered as a missed division or false negative. (Supplementary Fig. 3b). Finally, if a predicted parent node was identified as a division, but the daughters did not match the ground truth daughters, the division was counted as incorrect and included as a false negative division(Supplementary Fig. 3c). Finally, any remaining predicted divisions that cannot be matched to a ground truth division are counted as false positives.

We utilized the classified divisions to calculate a set of summary statistics, including recall, precision, and F1 score. Additionally, we utilized the mitotic branching correctness (MBC) metric defined by Ulicna *et al.*^30^, calculated as follows:

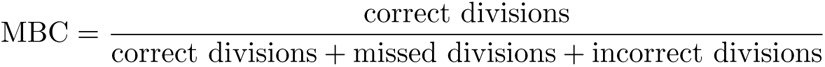

### 3.6 Deployment

We previously described the DeepCell Kiosk^55^, our scalable cloud-based deployment for deep learning models. The Kiosk provides a drag-and-drop interface for model predictions currently deployed at www.deepcell.org/predict. To provide a seamless pipeline for nuclear segmentation and tracking, we deployed a new consumer for tracking jobs. First, each movie is split into single frames, which are distributed for nuclear segmentation. This step takes advantage of the Kiosk’s ability to parallelize and scale resources to match demand. Once nuclear segmentation is complete on all frames, the masks are concatenated, and tracking is performed. The user receives a final output that contains the raw data, labeled masks, and lineages.

### 3.7 Benchmarking

We compared the performance of our model against four other algorithms: Baxter^51^, CellTrackerGNN^52^, EmbedTrack^53^, and Trac^x36^. Using the test split of our dataset, we evaluated the tracking performance of each algorithm on ground truth segmentation and predicted segmentations generated by either the algorithm or Caliban. We evaluated the resulting tracking predictions using our division evaluation pipeline and evaluation software from the Cell Tracking Challenge^45^. To evaluate the performance of Caliban on CTC movies, we manually annotated the two test movies from the Fluo-N2DL-HeLa dataset. To preserve the integrity of the challenge, we have not released our annotations but used them for evaluation in this paper. The notebooks used to generate benchmarks are available at https://github.com/vanvalenlab/Caliban-2024_Schwartz_et_al.

We evaluated Caliban’s inference speed using a single GPU (NVIDIA RTX A6000) and eight CPUs (AMD EPYC 7763 64-Core Processor). The inference time was split into four sections: segmentation inference, neighborhood encoder inference, tracking inference, and linear assignment. Inference was repeated three times for each movie in the test data split.

## Declarations

### Code Availability

The software used for data labeling is available at https://github.com/vanvalenlab/deepcell-label. The code used for model development, cell tracking, and model deployment are available at https://github.com/vanvalenlab/deepcell-tf, https://github.com/vanvalenlab/deepcell-tracking, and https://github.com/vanvalenlab/kiosk-console, respectively. Finally, the code for reproducing all models and figures included in this paper is available at https://github.com/vanvalenlab/Caliban-2024_Schwartz_et_al. All code is released under a modified Apache license and is free for non-commercial use.

### Data Availability

The DynamicNuclearNet dataset is available through deepcell.datasets (https://deepcell.readthedocs.io/en/master/ data-gallery/dynamicnuclearnet.html#sphx-glr-data-gallery-dynamicnuclearnet-py) for non-commercial use.

### Author Contributions

MS, EM, GM, and DVV conceived the project. MS, EM, and GM developed the HITL labeling methodology. MS, EM, EB, RD, WG, and DVV developed the deep learning cell-tracking methodology. MS and DVV developed the deep-learning nuclear segmentation methodology. GM, TD, EB, and WG developed DeepCell Label and adapted it to live-cell images. GM developed new labeling tools within DeepCell Label to accelerate labeling. MS and WG developed the data-versioning methodology with DVC. MS, EM, GM, EP, and several unnamed image labelers developed the DynamicNuclearNet. MS and DVV developed the benchmarking framework. WG and DVV oversaw software engineering for DeepCell Label and Caliban. MS and DVV wrote the manuscript, with input from all authors. DVV supervised the project.

## Acknowledgments

We thank Sam Cooper, Jan Funke, Benjamin Gallusser, Uri Manor, Alan Lowe, Paul Blainey, Iain Cheeseman, and Manuel Leonetti for useful conversations and insightful feedback. We thank Takamasa Kudo and Nicolas Quach for live cell imaging data. We utilized the HeLa cell line in this research. Henrietta Lacks, and the HeLa cell line established from her tumor cells without her knowledge or consent in 1951, has made significant contributions to scientific progress and advances in human health. We are grateful to Lacks, now deceased, and to the Lacks family for their contributions to biomedical research. This work was supported by awards from the Shurl and Kay Curci Foundation (to DVV), the Rita Allen Foundation (to DVV), the Susan E. Riley Foundation (to DVV), the Pew-Stewart Cancer Scholars program (to DVV), the Gordon and Betty Moore Foundation (to DVV), the Schmidt Academy for Software Engineering (to TD), the Michael J. Fox Foundation through the Aligning Science Across Parkinson’s consortium (to DVV), the Heritage Medical Research Institute (to DVV), and the National Institutes of Health New Innovator program (DP2-GM149556) (to DVV).

## Supplement

Supplementary File 1: DeepCell Label User Manual

Supplementary File 2: Segmentation Correction Instructions

Supplementary File 3: Tracking Correction Instructions

**Supplementary Table 1:** Benchmarking the performance of different tracking methods on the test split of DynamicNuclearNet. Bold font indicates the best scores on predicted segmentations. Italic font denotes the best scores on ground truth (GT) segmentations. CTC: Cell Tracking Challenge, DET: CTC detection accuracy^45^, SEG: CTC segmentation accuracy^80^, TRA: CTC tracking accuracy^80^, Div R: division recall, Div P: division precision, Div F1: division F1 score, MBC: mitotic branching correctness^30^, AA: association accuracy^46,47^, TE: target efficiency^46,47^.

| Tracking | Segmentation | DET | SEG | TRA | Div R | Div P | Div F1 | MBC | AA | TE |
| --- | --- | --- | --- | --- | --- | --- | --- | --- | --- | --- |
| Caliban | Caliban | **0.990** | **0.913** | **0.989** | **0.90** | **0.77** | **0.83** | **0.71** | 0.95 | 0.96 |
|  | GT | *1.000* | *1.000* | *0.999* | *0.95* | *0.92* | *0.94* | *0.88* | 0.98 | 0.98 |
| Baxter | Caliban | 0.988 | 0.908 | 0.986 | 0.49 | 0.73 | 0.59 | 0.41 | **0.98** | **0.98** |
|  | GT | 0.997 | 0.996 | 0.997 | 0.60 | 0.89 | 0.72 | 0.56 | *1.00* | *0.99* |
| Trac<sup>x</sup> | Caliban | **0.990** | **0.913** | **0.989** | 0.35 | 0.29 | 0.32 | 0.19 | 0.95 | 0.95 |
|  | GT | *1.000* | *1.000* | 0.999 | 0.34 | 0.54 | 0.42 | 0.27 | 0.98 | 0.98 |
| CellTrackerGNN | CellTrackerGNN | 0.815 | 0.682 | 0.812 | 0.43 | 0.05 | 0.08 | 0.04 | 0.76 | 0.76 |
|  | GT | *1.000* | 0.999 | 0.996 | 0.65 | 0.10 | 0.18 | 0.10 | 0.93 | 0.93 |
| EmbedTrack | EmbedTrack | 0.816 | 0.642 | 0.815 | 0.64 | 0.17 | 0.27 | 0.15 | 0.82 | 0.82 |

**Supplementary Table 2:** Benchmarking the performance of different tracking methods on the CTC Fluo-N2DL-HeLa test movies. Bold font indicates the best scores on predicted segmentations. Italic font denotes the best scores on ground truth (GT) segmentations. CTC: Cell Tracking Challenge, DET: CTC detection accuracy^45^, SEG: CTC segmentation accuracy^80^, TRA: CTC tracking accuracy^80^, Div R: division recall, Div P: division precision, Div F1: division F1 score, MBC: mitotic branching correctness^30^, AA: association accuracy^46,47^, TE: target efficiency^46,47^.

| Tracking | Segmentation | DET | SEG | TRA | Div R | Div P | Div F1 | MBC | AA | TE |
| --- | --- | --- | --- | --- | --- | --- | --- | --- | --- | --- |
| Caliban | Caliban | **0.994** | **0.897** | **0.993** | **0.77** | **0.88** | **0.82** | **0.70** | **0.99** | **0.99** |
|  | GT | *1.000* | *1.000* | *0.999* | 0.81 | *0.90* | *0.85* | *0.74* | *0.99* | *0.99* |
| Baxter | Caliban | 0.992 | 0.894 | 0.991 | 0.73 | 0.82 | 0.77 | 0.63 | **0.99** | **0.99** |
|  | GT | 0.994 | 0.993 | 0.993 | 0.74 | 0.81 | 0.78 | 0.63 | 0.98 | 0.98 |
| Trac<sup>x</sup> | Caliban | 0.994 | 0.896 | 0.991 | 0.09 | 0.44 | 0.15 | 0.08 | 0.98 | 0.98 |
|  | GT | *1.000* | *1.000* | 0.997 | 0.07 | 0.45 | 0.13 | 0.07 | 0.98 | 0.98 |
| CellTrackerGNN | CellTrackerGNN | 0.931 | 0.846 | 0.929 | 0.51 | 0.24 | 0.33 | 0.20 | 0.92 | 0.92 |
|  | GT | 0.998 | 0.996 | 0.996 | *0.90* | 0.38 | 0.53 | 0.36 | 0.96 | 0.96 |

**Supplementary Figure 1:**
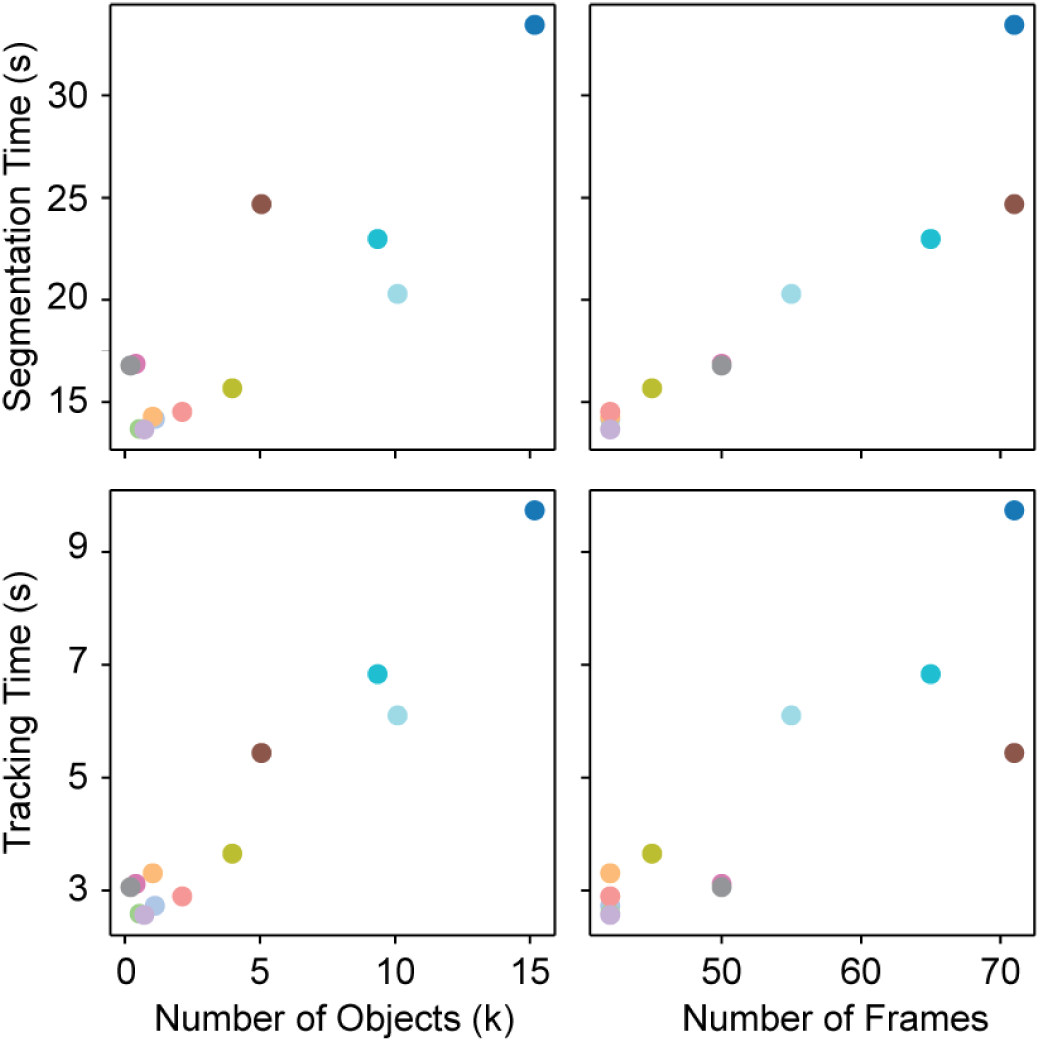
Runtime for segmentation and tracking with Caliban. The total runtime for segmentation and tracking is plotted as a function of the number of objects and frames in the sample. Each point represents a movie in the test data split, with a unique color assigned to each movie.

**Supplementary Figure 2:**
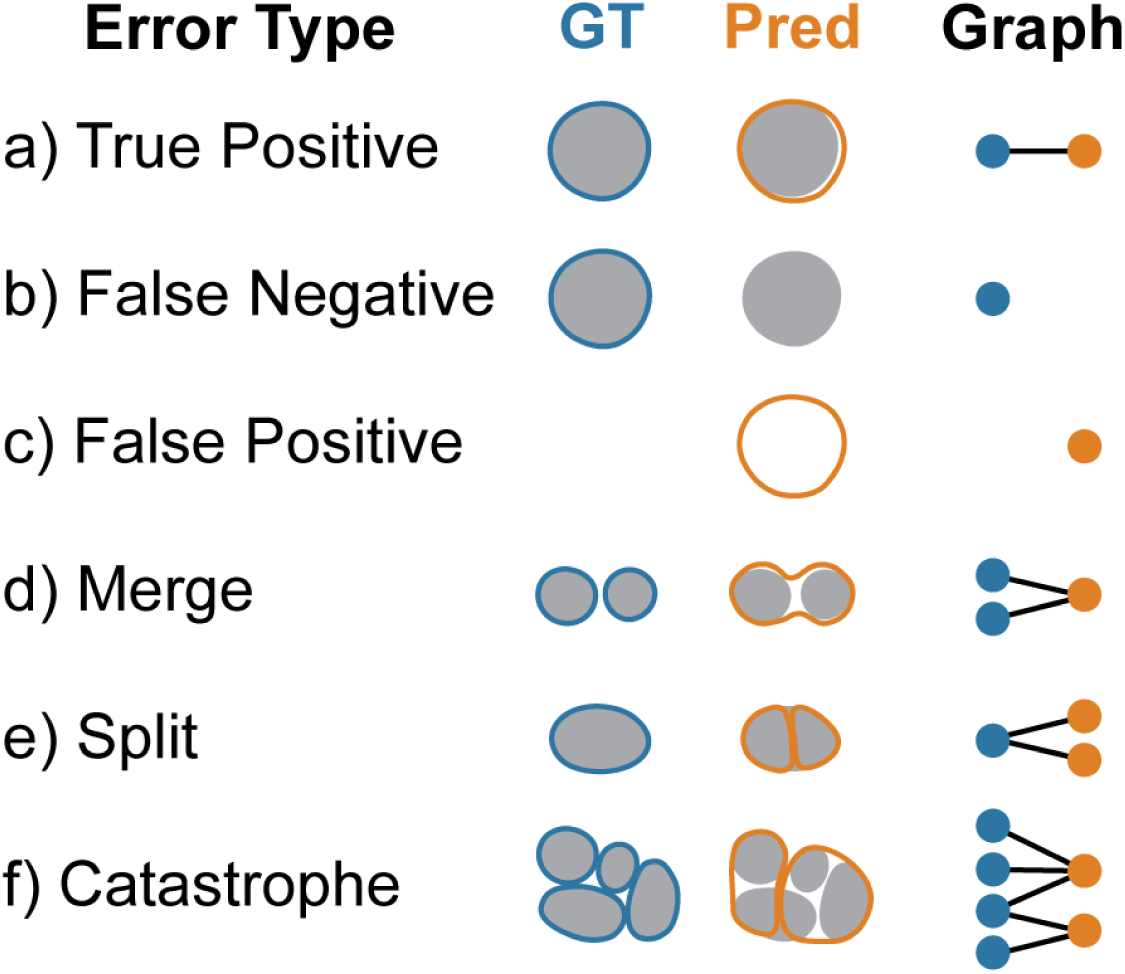
Object-based evaluation of segmentation performance. Segmentation predictions were evaluated based on object-level accuracy by first constructing a graph in which edges indicate an overlap between two objects. Each subgraph is then isolated and analyzed to identify the type of segmentation error present. (a) Subgraphs with one ground truth (GT) and one predicted node represent a true positive segmentation. Subgraphs containing only one node represent (b) a false negative if the node is ground truth or (c) a false positive if the node is predicted. Subgraphs with three nodes indicate (d) a merge if two ground truth nodes are associated with one predicted node or (e) a split if two predicted nodes are associated with one ground truth node. (f) Finally, all subgraphs containing more than three nodes are assigned to the catastrophe error class.

**Supplementary Figure 3:**
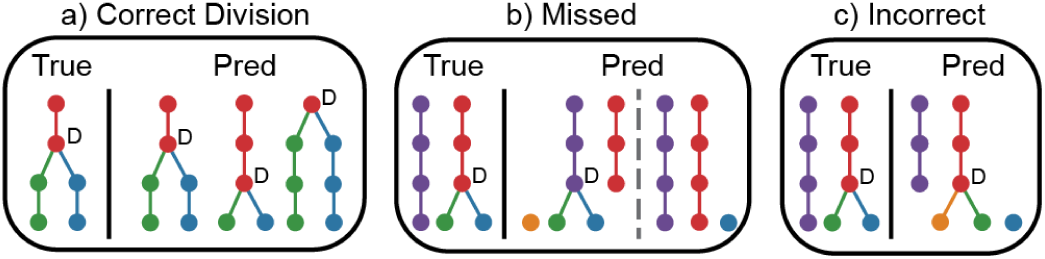
Division-based evaluation of tracking performance. Division events are classified as correct, missed, or incorrect based on a comparison of the true and predicted tracking graphs. (a) A division is considered correct if the prediction links the parent to the correct daughters within one frame of the ground truth division event. We allow divisions to shift in time because segmentation predictions can change when the cell is identified as one or two objects. (b) Divisions are identified as missed if the daughter cells are assigned to the incorrect parent or if no parent is identified. (c) A division is incorrect if the parent is assigned to only one of the correct daughter cells.

**Supplementary Figure 4:**
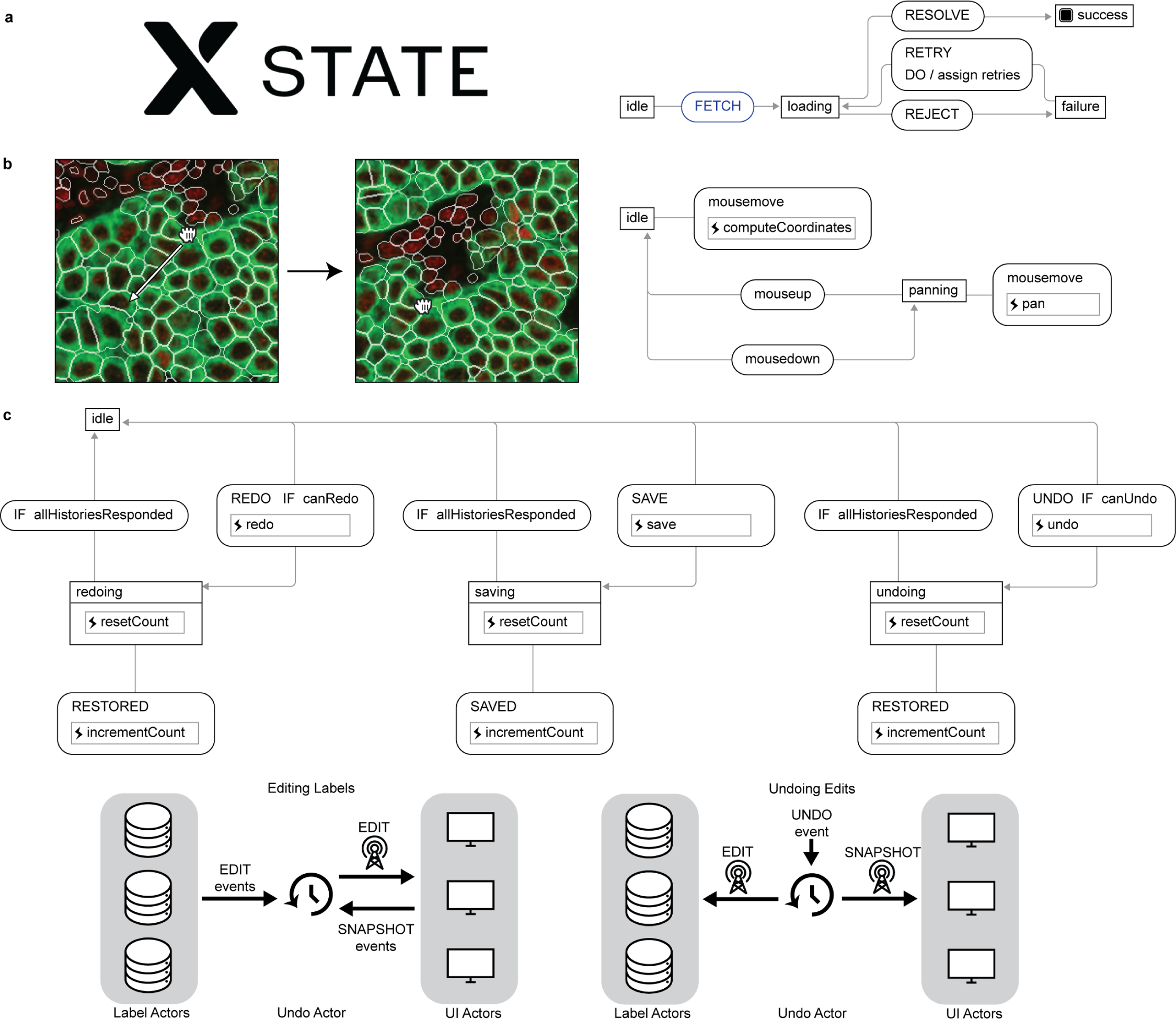
State management in the DeepCell Label backend. (a) DeepCell Label uses XState, a library for event-driven programming and state management, to drive its user interface (UI) and data labeling logic. Actors defined with XState control interactions with UI components and with labeled data. Actors consist of a context with arbitrary data, such as the settings for UI component or data labels; and a set of finite states and transitions between them. Actors receive events, which trigger transitions between states. Events are sent by user interactions or sent from other actors. (b) The panning states for the canvas actor. The actor begins in an idle state, where mousemove events update the position of the cursor. Upon a mousedown event, the actor transitions into a panning state, where mousemove events instead change the position of the image, enabling the user to browse the image. Once the user releases the mouse and triggers a mouseup event, the actor returns to the idle state. (c) The undo actor broadly orchestrates state across all UI and data labeling actors to enable undoing and redoing edits to labeled data. Actors that wish to subscribe to undo and redo events send an event to the undo actor to register itself as a UI actor or a data labeling actor. When data labeling actors edit their labels, they send a SNAPSHOT event to the undo actor, which then collects snapshots of all registered UI actors. When the user sends an undo event to the undo actor, the undo actor resends the snapshot events associated with the edit, globally restoring the application to its state just before the edit. Each registered actor maintains its own state and logic on how to implement undoable behavior, while the undo actor serves as a shared channel to coordinate registered actors.

