## Supplemental File 1 for "Caliban: Accurate cell tracking and lineage construction in live-cell imaging experiments with deep learning"

### Overview

In biology, cells naturally move over time. Occasionally, one cell divides into two cells. A cell that divides is a "parent cell" and the two new cells that result are "daughter cells." We want to track these movements and divisions. In this task, every cell in the movie has been labeled with a color and number. You will use DeepCell Label to correct the cell labels so that each cell has the same label across all frames. Additionally, you will find cell divisions and link the daughter cells to their parent cell.

---

### Your Task

For this job, each different label value is represented by a color and number. You will follow each cell throughout the movie and check that each cell:

- Has the same label value in every frame
- Is linked to its daughter cells if it divides
- If there is a division, that the two daughter cells have different labels from each other and its parent

DCL has tools that will help with tracking these cells. Each tool for each use case is discussed below.

Some labels may disappear or migrate out of the field of view. This is ok. You should **not** make new labels or edit the shape of these labels.

Check **each label in each frame** throughout the whole movie. **You must look at every cell label in each frame before submitting the job**.

### DCL tracking tools

When you click on the link you will see this (it may take a few moments to load):

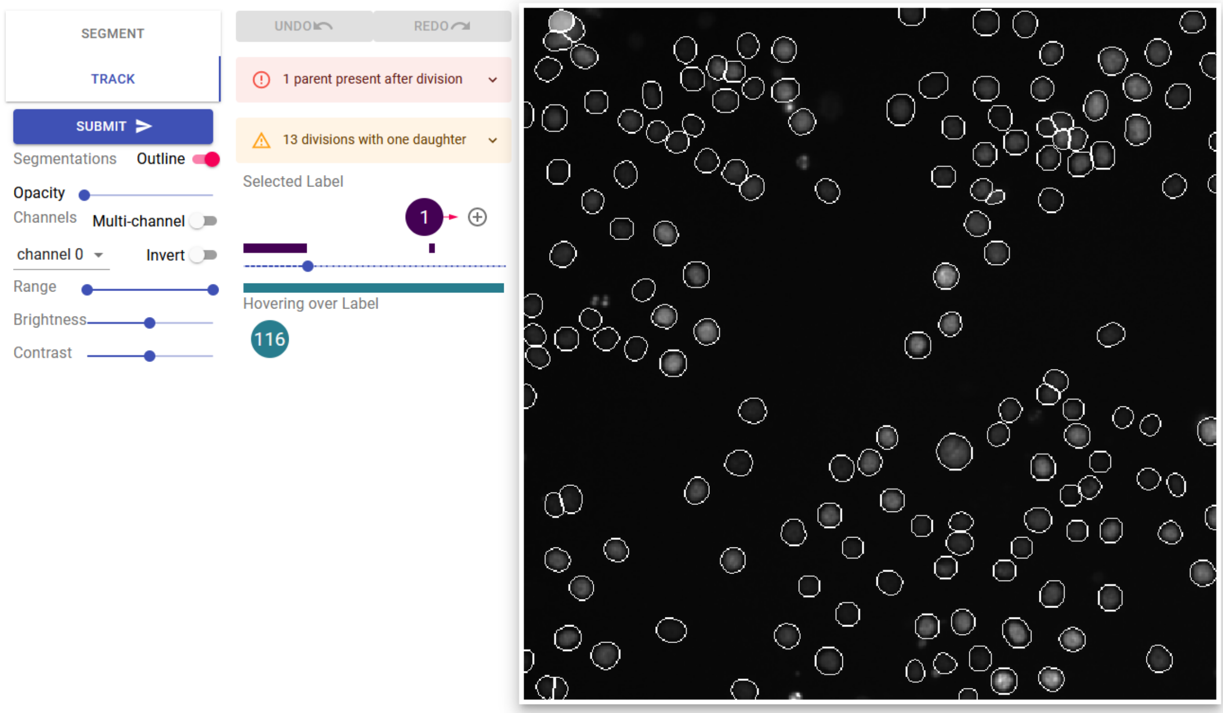


If you have used DCL before, this will look familiar. This is the work area you will use in each job.

The work area is divided up into 3 areas:

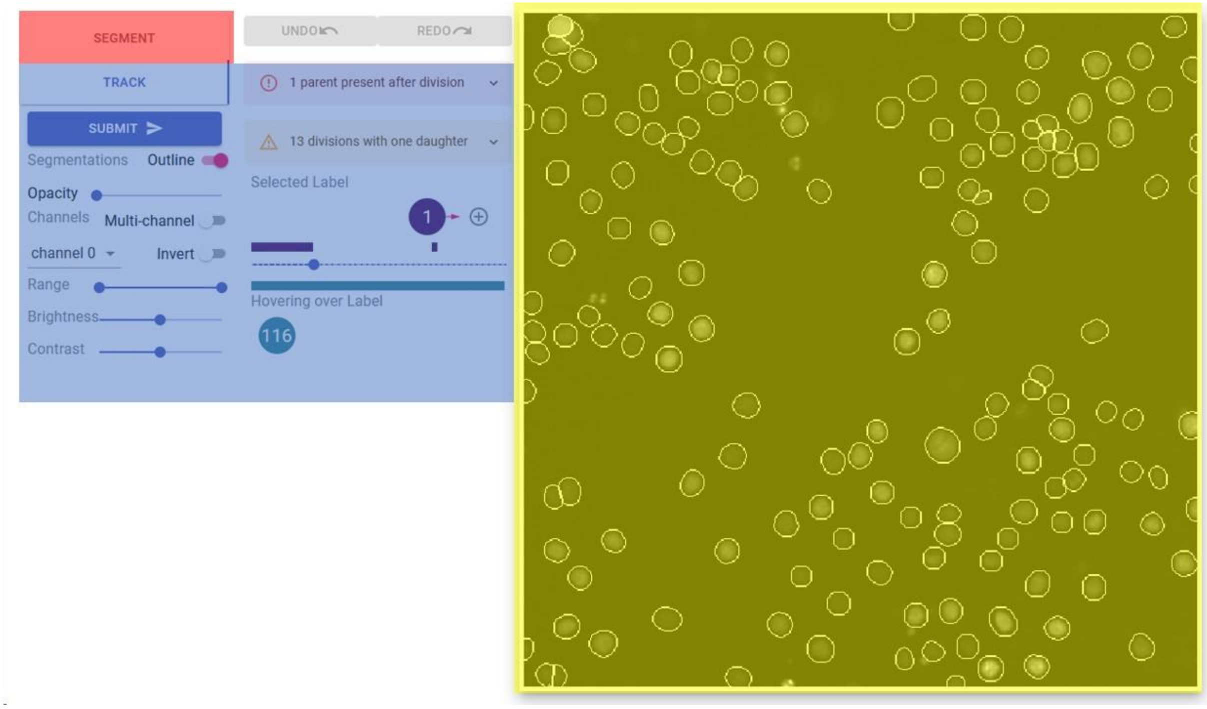


**Image Canvas (Yellow) -** This is where each frame of the movie and the cell labels will be displayed. The colors are there to help differentiate labels and to check if labels are staying the same throughout the movie.

**Annotation Tools (Red) -** These tools are for editing the label shapes or creating new ones. You should not need these tools for this task.

**Tracking Tools (Blue) -** These are the tools you will need to fix labels in the movie.

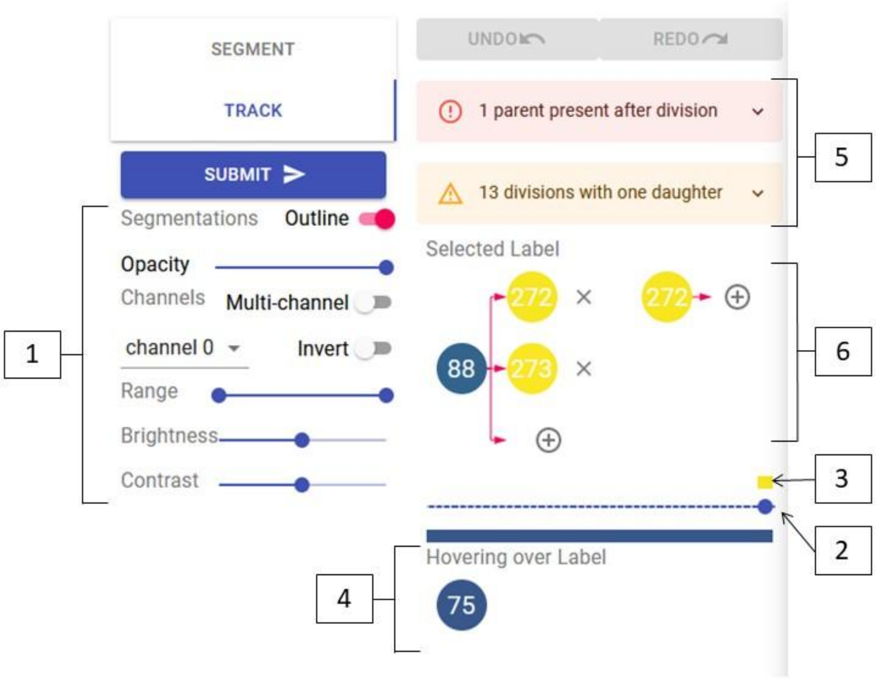


1. Image adjustments - These can help you see the movie better. For these jobs, increase the opacity to maximum so that only the labels are visible. You can press “Z” to change the opacity quickly.
2. Movie frame indicator - This shows which frame of the movie you are viewing. You can change frames by clicking on the bar or by pressing the “A” and “D” keys. “A” goes to the previous frame and “D” goes to the next frame.
3. Label frame indicator - This bar shows you in which frames the current selected label exists. (Note: the bar does not align perfectly to the movie frame indicator)
4. Cursor label indicator - This line shows the label value of the cell that the cursor is currently hovering over.
5. Error messages - The program checks for odd connections that exist in the movie. Pay close attention to these messages as they likely need to be corrected. Click on the messages to see labels that will need special attention. Red errors must be fixed. Yellow warnings need to be checked but may not require a change.
6. Lineage map - This map shows the currently selected cell and if that cell has a parent or daughters in the movie.

### Interactions with the image canvas

In this job, you will correct incorrect label assignments throughout the movie. Use the following keybinds to interact with the image canvas:

“A” - moves one frame backwards in time

“D” - moves one frame forward in time

“[“ - Selects the previous label value (values will loop from end to beginning)

“]” - Selects the next label value (values will loop from end to beginning)

“Z” - Increases the opacity of the labels

### General Instructions

The easiest way to interact with each job is to look at the label values to see when new cells appear and if labels change values in-between frames. New cells have a larger label number and will be yellow in color. These new cells are likely to be daughter cells as a result of cell division.

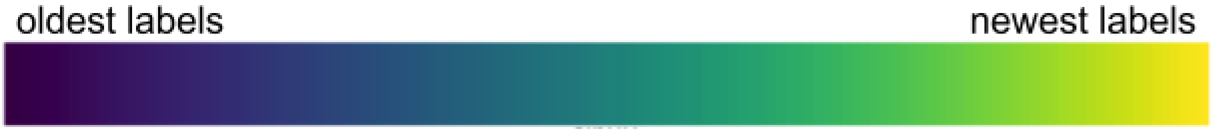


To set this up:

1. Press “Z” twice or increase the opacity slider to 100%
2. Press “[“ to select the highest numbered label
3. Click on the label number in the lineage map

This will take you to the last frame of this movie. You will then press “A” and “D” to cycle between frames in the movie. Follow the selected label through all the frames it is present in.

During this process, assess the following:

1. Does the label exist throughout every frame of the movie?
    a. YES: does the label stay constant throughout the movie?
        i. YES: move on to the next highest label by pressing “[“
        ii. NO: does the label switch places with another label?

            1. YES: see Swapping labels section
            2. NO: you likely have a situation where the parent label has been transferred to a daughter cell. See Changing parent cell to daughter cells section
    b. NO: move on to #2
2. Does the label disappear because the cell merges with another cell?
    a. YES: Is the current label a daughter of the cell it merges into?
        i. YES: Is the other daughter cell also associated with the parent cell.
            1. YES: move on to the next highest label by pressing "[“
            2. NO: See Changing parent cell to daughter cells or Adding daughter cells section
        ii. NO: see Adding daughter cells section
    b. NO: Has the label of that cell changed in a previous frame?
        i. YES see Combining labels section
        ii. NO: move on to #3
3. Does the cell disappear off the canvas?
    a. YES: scan all frames to see if the label stays the same if the cell reappears in the frame. Does the label stay the same?
        i. YES: move on to the next highest label by pressing "[“
        ii. NO: see Combining labels section and Swapping labels section
    b. NO: the label has disappeared due to cell death. Move on to the next highest label by pressing "[“

These general instructions should guide you through 90% of a job. Sometimes a cell needs multiple actions to correct it's label. Pay closer attention when correcting those cells. In the hardest cases, you may need to change a label to a new label value to correct all the cells. In that case, please see the Add new labels section

When you have finished reviewing all the labels, press the Submit button to complete your job.

### Adding daughter cells

Use these instructions to add daughter cells to a parent cell when a cell divides.

1. Select the parent cell
2. Click the "+" symbol next to the parent cell label in the lineage map and select Add Daughter from the menu

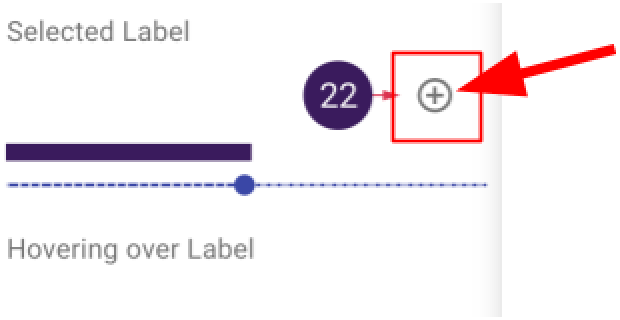


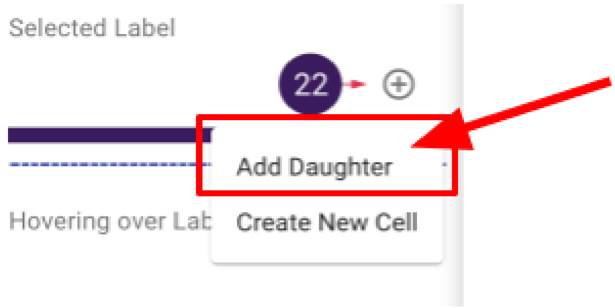


3. Scan through the movie until the daughter cells appear (usually the next frame of the movie after the parent label ends)
4. Click on a daughter cell on the canvas

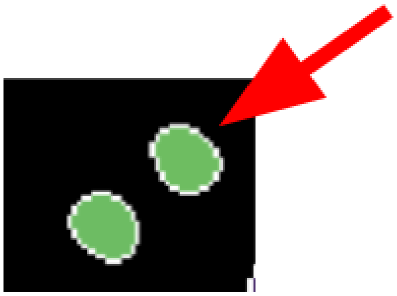

5. Wait for the lineage map to update

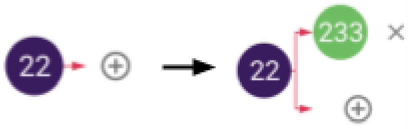


6. Repeat for the other daughter cell if necessary

#### When a parent label should be a daughter

Use these instructions when a parent cell divides but does not have a different label after dividing. A correct division will have different labels for the parent and daughters.

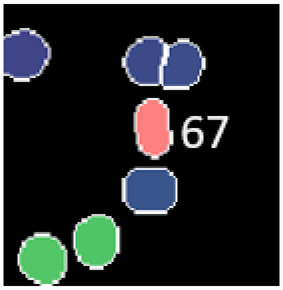


Frame 33

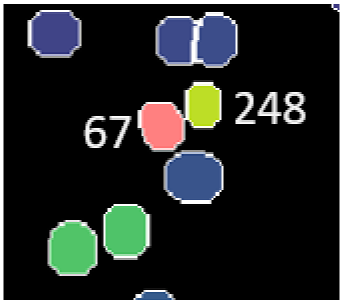


Frame 34

Example: Label 67 divides into two daughter cells in frame 34. Label 67 in frame 34 should be a new label instead of keeping the same label as before the division.

1. Select the parent cell

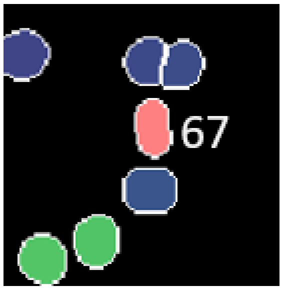


Frame 33

2. Click the "+" symbol next to the parent cell label in the lineage map and pick **Add Daughter** from the menu

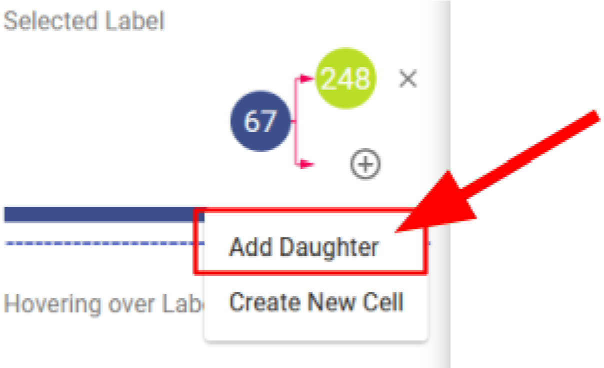


3. Scan to the frame where the parent cell divides

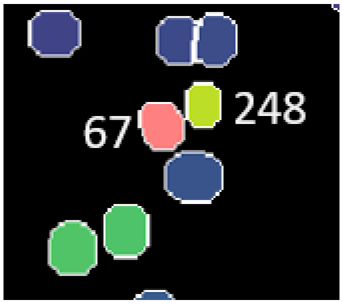


Frame 34

4. Click on the parent label in the canvas in the first frame where it becomes a daughter
5. Wait for the lineage map to update

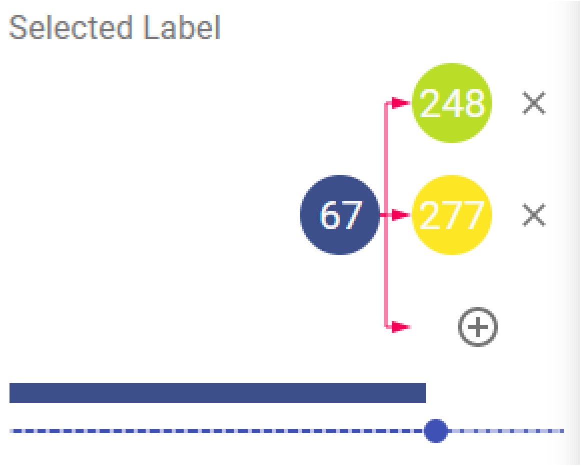


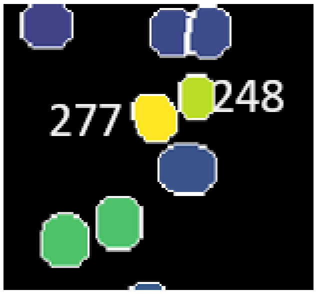


Frame 34

The daughter will have a new label  from this frame on.

#### Combining labels across frames

Use these instructions when one cell has different labels in different frames but should have the same label. Pay attention to these instructions as they are currently not easy.

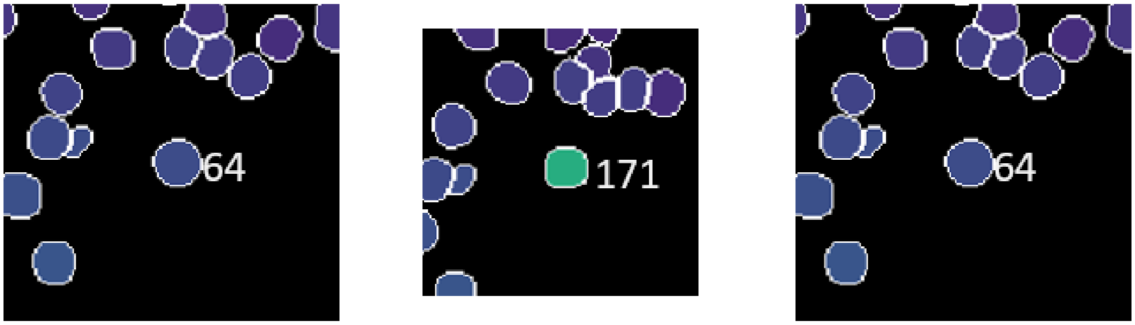


Frame 6 Frame 7 Frame 9

Example: Label 64 switches to 171 in frame 7.

1. Select one of the labels to be combined (usually the lower number)

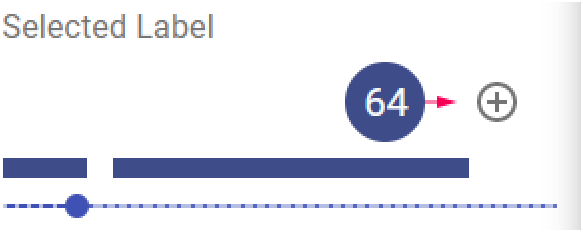


2. Click the “+” symbol next to the parent cell label in the lineage map and select “Add Daughter”

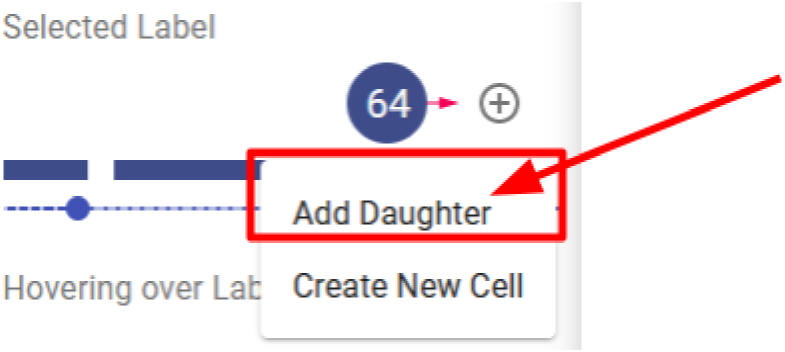


3. Scan to the frame where the cell changes to a different label without dividing (make sure the first label does not exist in this frame) and click on the new, different label you want to replace.

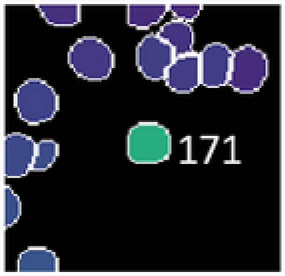


Frame 7

4. Wait for the lineage map to update

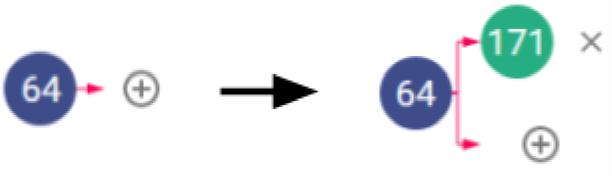


5. Click on the “x” symbol to the right of the daughter label and select “Replace with Parent” from the menu

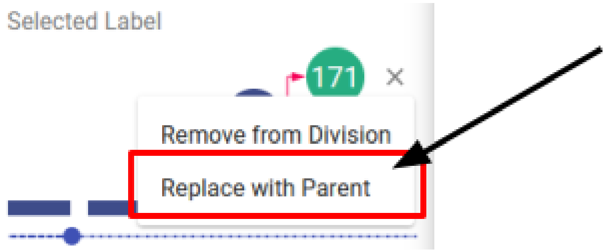


6. Wait for the lineage map to update

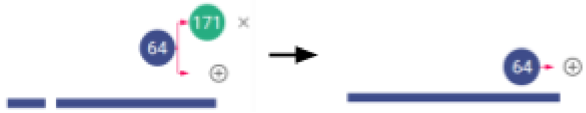


#### Swapping labels

Use these instructions to switch label values between two different labels.

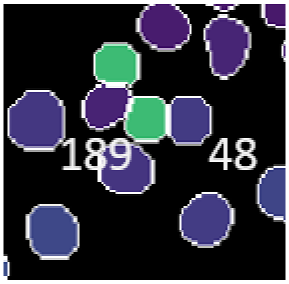


Frame 20

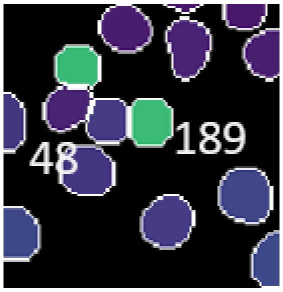


Frame 21

Example: Labels 189 and 48 switch places in frame 21.

1. Click the Segment button to change tools.

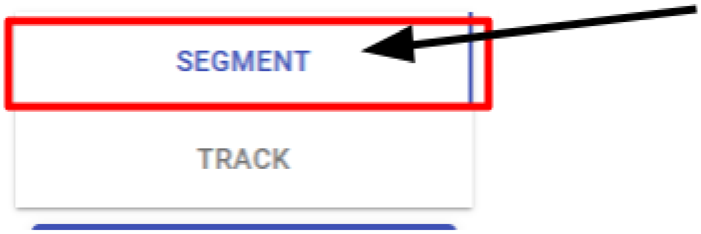


2. Make sure you are using the Select tool under the Tools buttons

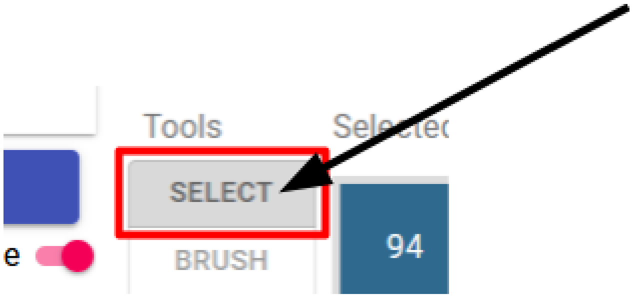


3. Double click on one of the labels to be changed so it is outlined in red

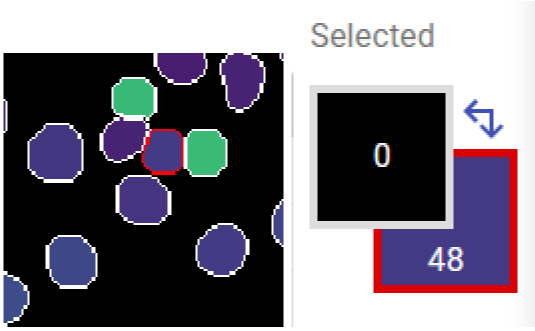


4. Click on the other label so it becomes red inside

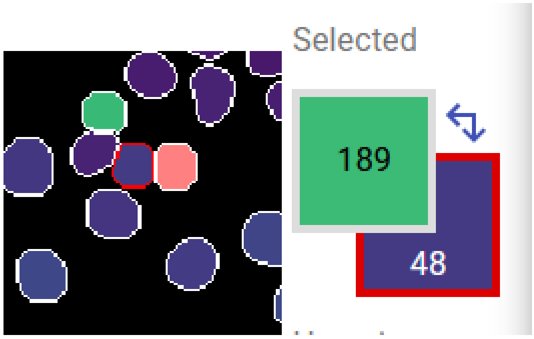


5. Click “Swap” under Actions

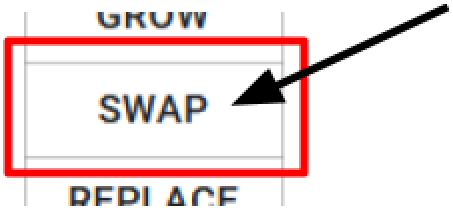


6. Wait for the highlighted cell to switch spots

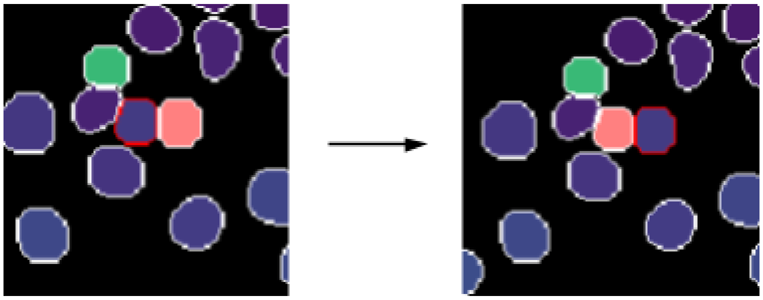

7. (Optional) If cells in multiple frames need to be changed, press “D” to go to the next frame and click “Swap” for each frame
8. Press “Esc” to unselect the 2nd label

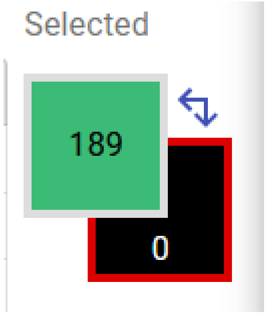


#### Add new cell value

Use these instructions to assign a new label value to cells that change labels multiple times between multiple cells during the course of the movie. **Use this function only when other functions will introduce a duplicate label in the same frame**.

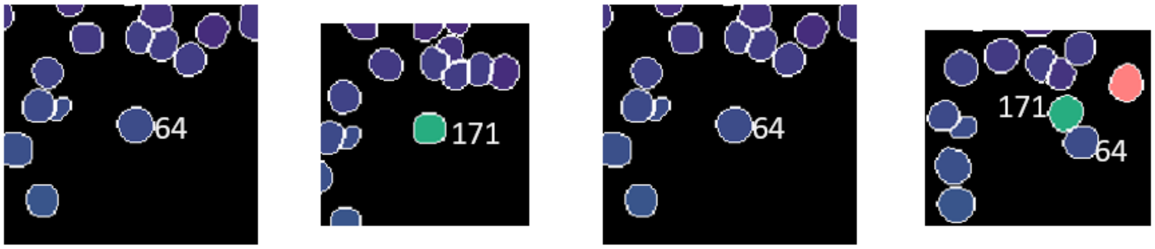


Frame 6 Frame 7 Frame 9 Frame 19

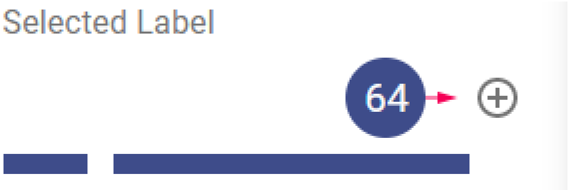


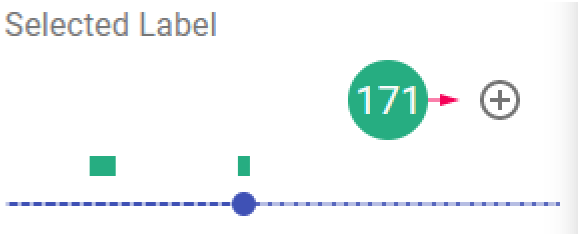


Example: Label 64 changes to 171 in frame 7 then changes back to 64 in frame 9. We cannot just combine label 64 with 171 because they both exist in frame 19. Label 171 in frame 19 will need to be changed to a new label before other fixes can occur.

Select the label that needs to be fixed

1. Advance the movie to the frame where that label reappears in the movie

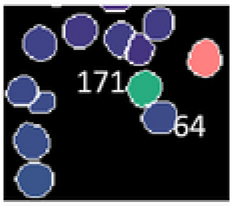


Frame 19

2. Click the "+" symbol next to the label in the lineage map and select "Create New Label"

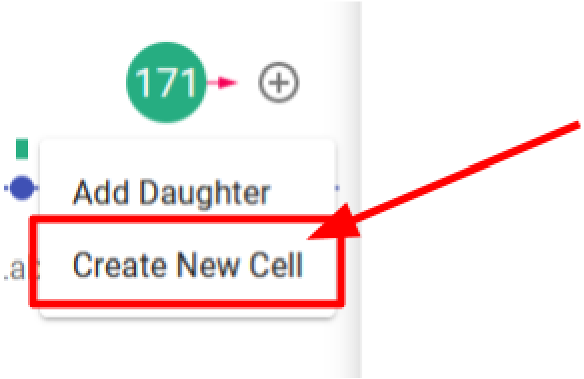


3. Wait for the label to change

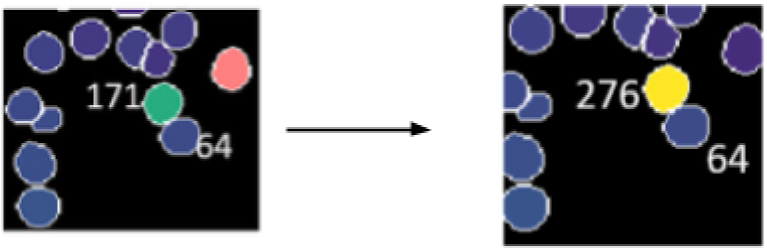


The new label replaces the original label from the current frame to the end of the movie. Then go back and make necessary corrections.

### If DCL doesn't work properly

First, make sure you have clicked inside the viewing window, which may solve your issue. This is a new tool that, despite testing, may still have some bugs. The tool may also have been updated since the last time you accessed it. If it does not seem to load correctly, try clearing your browser cache and refreshing the page. If this does not work and you encounter an error that prevents you from finishing correcting the file, please notify your manager and continue onto another file if possible. If you have any suggestions or questions, please share them with us! We would love to know what will make this tool easier and more useful for you.
