## Supplemental File 2 for "Caliban: Accurate cell tracking and lineage construction in live-cell imaging experiments with deep learning"

### How to Use DeepCell Label

This guide shows how to use different annotation tools within DeepCell Label (DCL) to create or edit annotated biological images. Please refer to job instructions for specific corrections needed. Use the outline as a reference for particular tool descriptions.

---

#### Overview

DCL offers a suite of tools to interact with labeled or unlabeled images. Unique project URLs should have already been generated for your project. If you need access to these URLs or want to generate your own, please contact a member of DeepCell for further information.

#### Layout

When you click on a valid DCL URL, you should be taken to a screen similar to this:

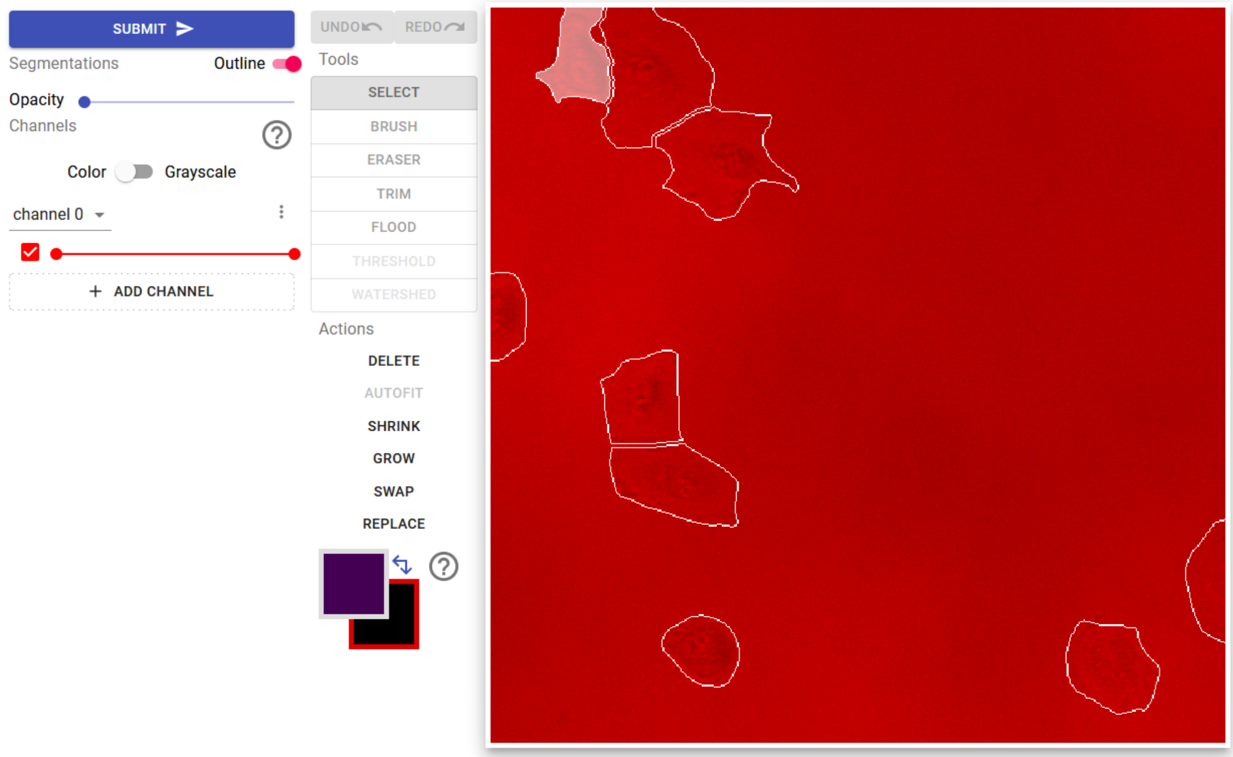

The right side of the screen is the workspace, which contains the image and any annotations associated with the file that has been uploaded. The left hand side contains all of the tools available to manipulate the image or annotations.

The **Undo/Redo Buttons** allows you to undo (or redo) the changes you make on the canvas. Be aware that switching tools and changing annotation selections are considered “edits” and multiple clicks of the button may be required to fully restore the desired result.

The **Palette Boxes** (located below the REPLACE button) allows you to visualize current label selections. The upper-left palette box shows the color of the currently selected label. Hovering over this box allows you to cycle through the different labels by clicking the **left and right arrows**, add a new label value by pressing the **“+” symbol**, or deselect the current label by pressing the **“x” symbol**. The lower-right palette box displays the label value of the canvas (black = no label value). The canvas is where edits can be made with tools and actions. The label value of the canvas can be changed to match any label (for more information, see the description of the **Select Tool**).

When you’ve finished your corrections, use the **Submit Button** to upload them.

#### Preparing the canvas:

When an image is loaded into DCL, the default view assumes your image contains multiple channels and will load your image with false coloring. Move the slider under each color to change the intensity of that color.

If you are working on phase images or single color images, you may want to switch the toggle to **Grayscale**. When in grayscale mode you are able to have more control over the brightness and contrast of the image. Move each slider to a position that allows you to see the most features within an image. These settings can be changed at any time.

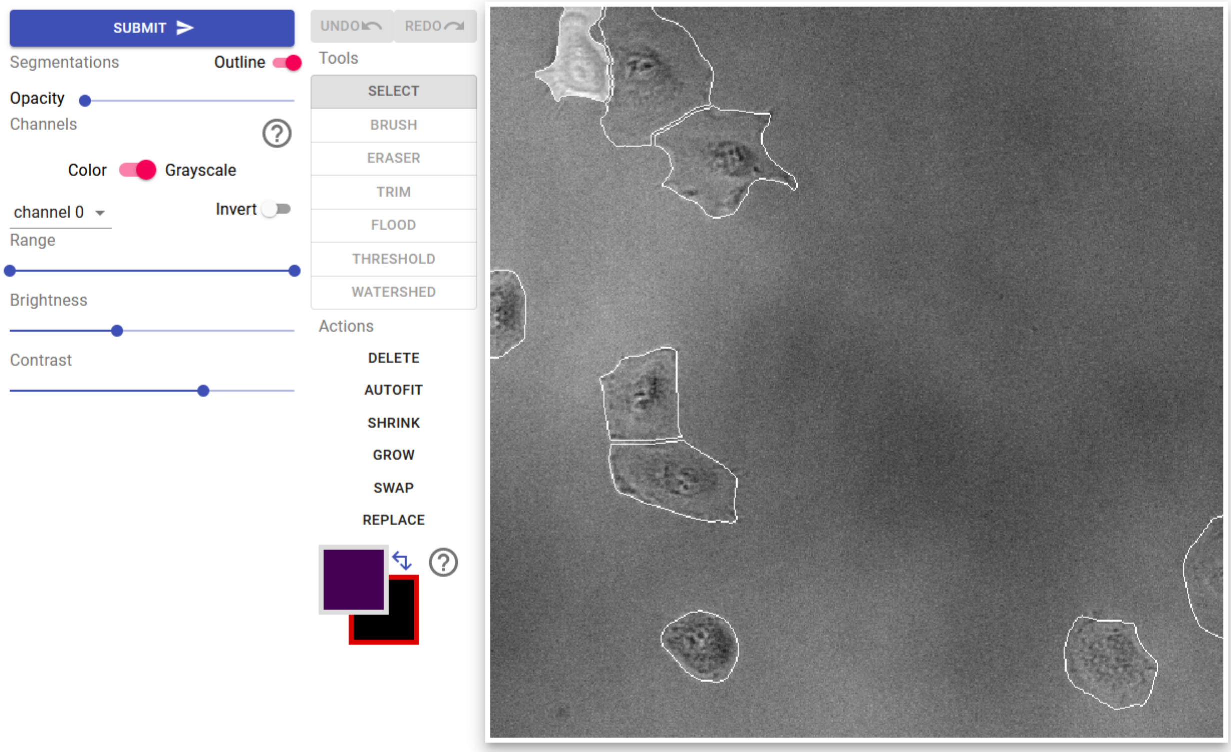


The outlines of any existing annotations will be displayed by default. They can be turned on/off by toggling the **Outline** button. If the image being annotated contains too many objects, it will be beneficial to see the annotations filled in with different colors. This can be done using the **Opacity** slider. Pressing “z” will also cycle through three preset opacity values (0, 30%, and 100%)

#### Zooming and panning:

To zoom into the image, use the scroll wheel or the “+” and “-” keys. To move around once zoomed in, hold down the spacebar then click and drag the mouse.

#### Changing channels:

Change channels with the "c" key.

##### Changing frames:

Use “a” or the left-arrow key to go back one frame, and “d” or the right-arrow key to go forward one frame. Going forward from the last frame will loop back around to the start of the file. Some datasets will only have one frame per file.

### Editing Tools

##### Select Tool:

Use the **select tool** to click on the annotation that you want to edit. Selected annotations will be highlighted in white and the annotation color shown in the upper-left palette box. Only selected cells can be interacted with through other tools or actions. Press “esc” to deselect the current annotation. If an object needs a new label, click the “+” symbols on the upper palette or press “n” to select an unused label. To select 2 labels at once (in the case of swapping or replacing), click on the label to replace twice (this will now be the background label in the lower-right palette box) and click on the label to paint with once (this will become the foreground label in the upper-left palette box).

##### Brush Tool:

Use the **brush tool** to make corrections to the current selected annotation. Click once to paint an area the size of the cursor. Click and drag to make larger edits. The brush size can be adjusted using the up- and down-arrow keys.

To adjust the border between 2 labels, first **select** the label you want to paint with, then hold down shift and click on the label you want to overwrite. Once you see the labels properly represented in the palette box (the top-left box will be the label you are painting with and the bottom-right box will be the label that is being brushed over), use the brush tool as you would normally.

##### Eraser Tool:

Use the **eraser tool** to remove portions of the current selected annotation. Click once to erase a portion of the label according to the size of the cursor. Click and drag to make larger edits. The eraser size can be adjusted using the up and down arrow keys. Press “z” to quickly switch to the **brush tool**.

##### Trim Tool:

Most of the time annotations should be connected. To quickly remove disconnected portions of an annotation, select the **trim tool** and click on the part of the annotation you want to keep. If the annotation to be fixed was not originally selected, click again for the trim tool to be applied.

##### Flood Tool:

Occasionally, annotations will contain empty spaces. To quickly fill these spaces, select the **flood tool** and click on the empty space to be filled. Make sure the correct annotation has been selected otherwise another label could be applied to the space.

#### Actions

Actions have been designed to make certain common edits easier to perform. Make sure to use the **select tool** to click on the annotation that needs to be corrected.

##### Delete:

To delete a whole annotation, click on that label with the **select tool** and the click **delete**. This is useful when whole objects are labeled incorrectly.

##### Shrink/Grow:

Annotations that are uniformly too large or too small can be edited using the **shrink** or **grow** actions. Both shrink and grow add or subtract one pixel around the label respectively.

##### Replace:

Sometimes annotations of a single cell have been split in error and given two different labels. To correct this, double click one of the labels, click the second label and click **replace.** Repeat as necessary. Use the **brush tool** to make all parts of the label connect.

##### Swap two labels:

This action is only needed to make sure that labels match up with each other correctly across different frames of a file. Select two different labels by double clicking on one label and then clicking on the other. Click **swap** to swap values.
