## Supplemental File 3 for "Caliban: Accurate cell tracking and lineage construction in live-cell imaging experiments with deep learning"

**DeepCell Label has been updated to improve the interface, so some functions and tools have changed. Please read the updated manual (DCL User Manual) for general instructions as to how to use this new tool. Once you are familiar with the new layout and features, refer to the following examples to check for specific work you need to do to complete the job. Thank you!**

### Overview

In this task you will work with a tool to create annotations of densely packed cells. This tool, DeepCell Label, has many features in it to draw and correct annotations. In this task, you will use a set of those features to create annotations of all the cells in the image.

### Background

Our group is in the process of writing a computer program to automatically identify cells in culture. To help us do this, we're asking you to help create annotated datasets where single cells are manually identified. We can then feed this into the computer program to teach it how to accurately identify cells by itself. This program will be used in other research laboratories to study a range of topics, including viruses, cancer cells, and immune cells. **The software we create is only as good as the data used to create it, so accuracy in your annotations is extremely important.**

In this job, each file contains many cells. Many of the cells are already labeled correctly, but some cells in the file have labels that are too big, too small, or overlap adjacent cells. You will correct the labels with the provided tools.

### Your Task

For this job, **your goal is to make sure that the border around each cell is in precisely the correct spot.**

### Algorithm Accuracy: High

This means that **<u>most</u>** of the cells in this image are correct. **<u>Some</u>** of the cells in this job will need to be changed.

### New Function: Autofit

A new function has been added to the tool set that automatically fills in missing labels.

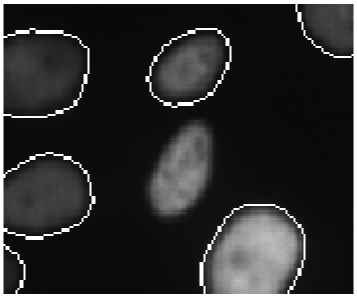

To use this function do the following:

1. Click on the brush tool

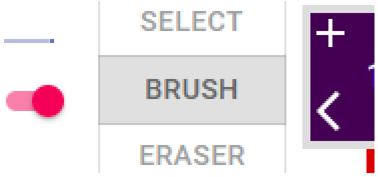


2. Press "N" to select a new label
3. Fill in a small portion of the cell to be labeled

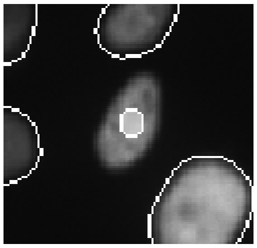

4. Click on Autofit

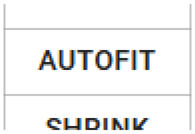


5. Wait until the label gets created

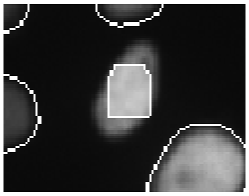

6. Evaluate the label and made edits as necessary

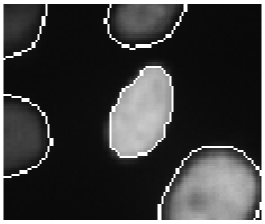

#### Specific Instructions for this job

The most common errors in this job are multiple cells with a single label. These labels need to be separated.

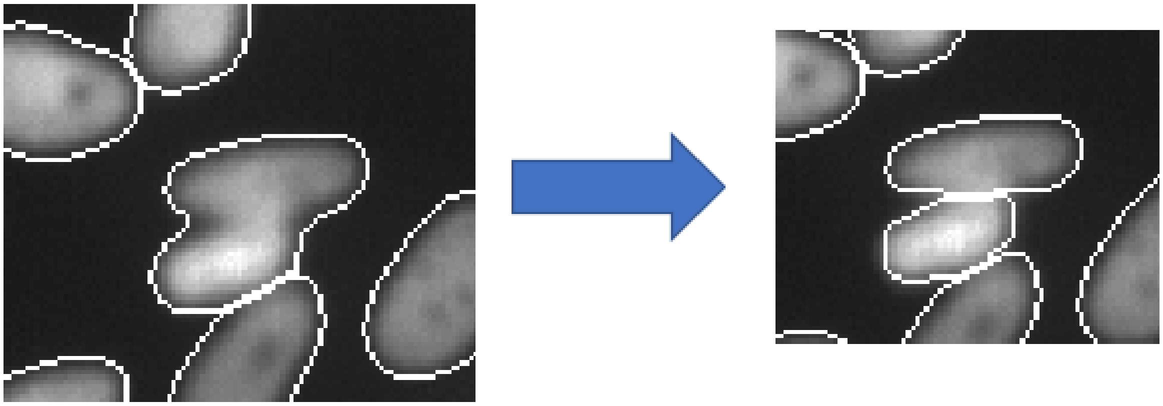

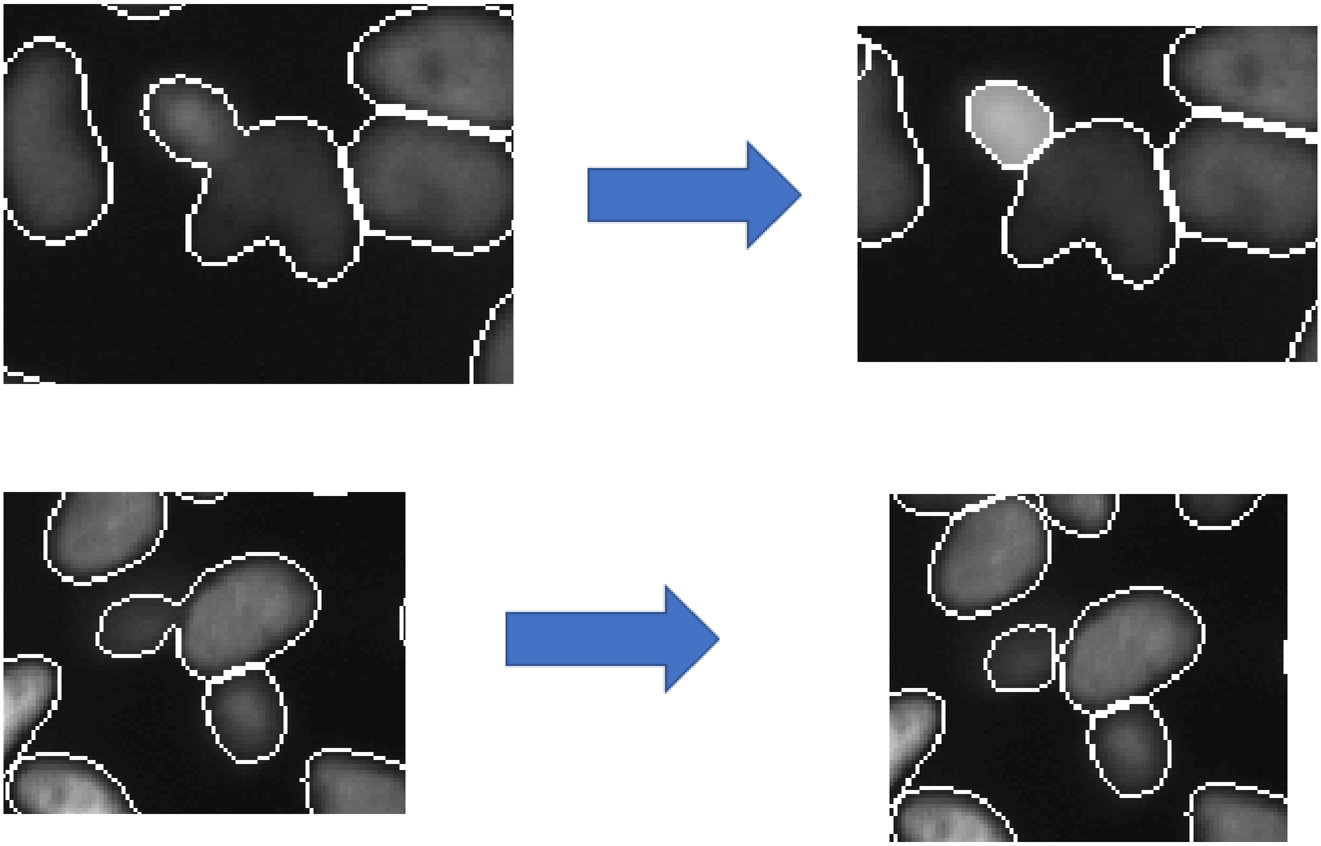

### General Instructions

Less common errors may also be found in this job. Watch out for and correct the following errors.

#### Label too large

When a cell has a label larger than the cell, fix the label to match the size of the cell.

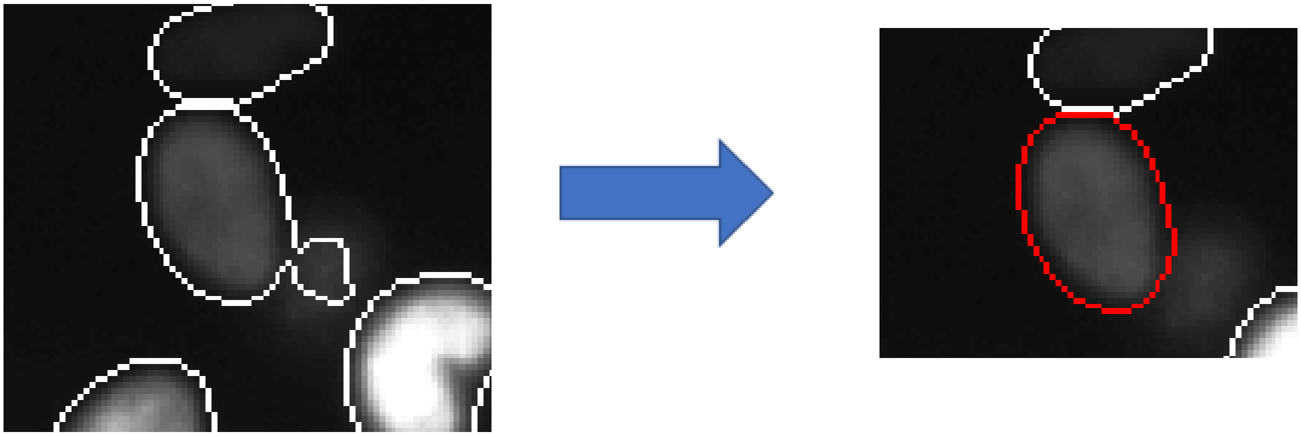

#### Missing cell

If a cell doesn't have a label, you should add one

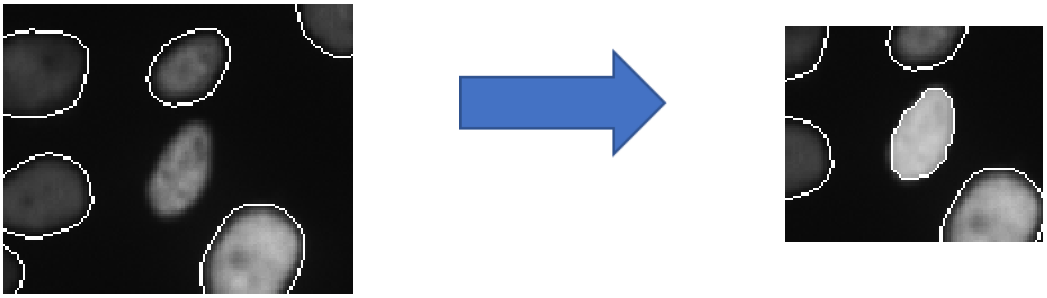

### If DCL doesn't work properly

First, make sure you have clicked inside the viewing window, which may solve your issue. This is a new tool that, despite testing, may still have some bugs. The tool may also have been updated since the last time you accessed it. If it does not seem to load correctly, try refreshing the page without cached data. If this does not work and you encounter an error that prevents you from finishing correcting the file, please notify your manager and continue onto another file if possible.
